# Yeast ATM and ATR use different mechanisms to spread histone H2A phosphorylation around a DNA double-strand break

**DOI:** 10.1101/2019.12.17.877266

**Authors:** Kevin Li, Gabriel Bronk, Jane Kondev, James E. Haber

## Abstract

One of the hallmarks of DNA damage is the rapid spreading of phosphorylated histone H2A (γ-H2AX) around a DNA double-strand break (DSB). In the budding yeast *S. cerevisiae*, nearly all H2A isoforms can be phosphorylated, either by Mec1^ATR^ or Tel1^ATM^ checkpoint kinases. We induced a site-specific DSB with HO endonuclease at the *MAT* locus on chromosome III and monitored the formation of γ-H2AX by ChIP-qPCR in order to uncover the mechanisms by which Mec1^ATR^ and Tel1^ATM^ propagate histone modifications across chromatin. With either kinase, γ-H2AX spreads as far as ∼50 kb on both sides of the lesion within 1 h; but the kinetics and distribution of modification around the DSB are significantly different. The total accumulation of phosphorylation is reduced by about half when either of the two H2A genes is mutated to the nonphosphorylatable S129A allele. Mec1 activity is limited by the abundance of its ATRIP partner, Ddc2. Moreover, Mec1 is more efficient than Tel1 at phosphorylating chromatin in *trans* – at distant undamaged sites that are brought into physical proximity to the DSB. We compared experimental data to mathematical models of spreading mechanisms to determine whether the kinases search for target nucleosomes by primarily moving in three dimensions through the nucleoplasm or in one dimension along the chromatin. Bayesian model selection indicates that Mec1 primarily uses a 3D diffusive mechanism, whereas Tel1 undergoes directed motion along the chromatin.

## Introduction

Genetic loci can be separated by thousands to millions of base pairs in the nucleus, constraining contact between distant regions of the genome. Yet many biological processes rely on the effective communication between distant parts of the genome to facilitate a diverse range of phenomena such as the regulation of gene expression through promoter-enhancer interactions, the formation of chromatin loops during chromosome condensation or the initiation of homologous recombination in the presence of DNA breaks (1–3). Therefore, an understanding of many nuclear processes can be uncovered by investigating how distant chromosomal regions can establish genomic interactions in a timely fashion (4).

Communication between remote parts of the genome is often facilitated by the presence of intermediary proteins that shuttle information in a three-dimensional (3D) manner through the nucleoplasm or in a one-dimensional (1D) manner along the chromatin. 3D modes of communication lead to both intra- and interchromosomal interactions either by the physical folding of chromatin to bring two regions of the genome within close proximity (looping) or by 3D diffusion of a protein from one genetic locus to another through the surrounding nucleoplasm (5). In both looping and 3D diffusion, information can be transferred over many kilobases without having to interact with the intervening chromatin. In contrast, 1D mechanisms are intrinsically intrachromosomal and communication is predicated on the movement of DNA-bound proteins along the contours of the chromatin fiber. 1D mechanisms include either diffusive or unidirectional motion of proteins along the chromatin (5). Comparisons of experimental data with biophysical models of chromatin looping, 3D diffusion, 1D diffusion and directed sliding provides insight into how specific information transfers take place in the nucleus.

In particular, we are interested in studying how genomic interactions are achieved over a distance of many kilobase pairs (kb) after DNA damage. The rapid formation of γ-H2AX, the phosphorylated form of histone H2A (H2AX in mammals), over an extensive region of the chromatin is one indicator of DNA double strand breaks (DSBs) (6–9). In budding yeast, γ-H2AX is formed by the phosphorylation at H2A-S129 by the checkpoint kinases Mec1 and Tel1, homologs of mammalian ATR and ATM, respectively. Previous studies have shown that Mec1 and Tel1 are both capable of forming γ-H2AX regions up to 50 kb on either side of a DSB (9), while their mammalian homologs, especially ATM, are able to phosphorylate histones in chromatin regions in excess of 1 Mb in mammalian cells (6, 7). γ-H2AX has been shown to play a role in the recruitment and retention of factors responsible for efficient DNA repair, DNA damage signaling and chromatin remodeling, but is not essential for these processes (8, 9).

Mec1 and Tel1 are activated in the presence of a DSB by different mechanisms. In budding yeast, as in mammals, Tel1^ATM^ is attracted to a broken chromosome end by its association with Mre11-Rad50-Xrs2^Nbs1^ (13). In contrast, the recruitment of Mec1^ATR^ requires that the broken end undergoes some 5’to 3’ resection, promoting the binding of the single-strand DNA-binding protein complex, RPA. RPA is then bound by Ddc2^ATRIP^, which is the obligate partner of Mec1^ATR^ (14, 15). While we know that both Mec1 and Tel1 are actively recruited to DSBs and are involved in γ-H2AX formation, the means by which Mec1 and Tel1 kinases reach histones tens of kilobases from the DSB is still unknown. To address this problem, we monitored the kinetics and extent of γ-H2AX formation in budding yeast, by ChIP-qPCR, after creating a synchronously-induced DSB at a specific location on chromosome III. We compared experimentally measured γ-H2AX distributions to mathematical formulations of different phosphorylation spreading mechanisms to determine whether the kinases can reach histones far from the DSB using chromatin looping, 3D diffusive, 1D diffusive or directed sliding mechanisms. Through Bayesian model selection, we conclude that Mec1 reaches target histones by a 3D diffusive mechanism, while Tel1 dynamics is best described by directed 1D sliding along the chromatin.

## Results

### Mec1 and Tel1 kinases use different mechanisms for γ-H2AX spreading

We studied phosphorylation spreading in *S. cerevisiae* after inducing a site-specific DSB using a galactose-inducible HO endonuclease, resulting in robust cleavage at the *MAT* locus on chromosome III within 30 min (Figure S1). DSBs were rendered irreparable by homologous recombination through the deletion of both *HML* and *HMR* donor loci (16). γ-H2AX spreading was studied in G1-arrested haploid *MAT***a** cells where the re-ligation of cleaved ends by nonhomologous end-joining was prevented by deleting *NEJ1* or *YKU80* (17). The formation of γ-H2AX over a region of roughly 50 kb on both sides of the DSB was quantified by ChIP-qPCR with an antibody specific to phosphorylated H2A-S129. To differentiate between the mechanisms used by Mec1 and Tel1 to phosphorylate large regions of chromatin adjacent to the break, we measured γ-H2AX levels in strains where both Mec1 and Tel1 were active or when only Mec1 or Tel1 kinase was active.

In G1-arrested cells, 5’ to 3’ resection of DSB ends is blocked (12, 13). Under these conditions, the absence of RPA binding leads to lack of Mec1 recruitment; so only Tel1 is active (9). We confirmed that in *nej1*Δ G1-arrested cells, Tel1 robustly modifies the chromatin adjacent to *MAT*, but there is only a very small signal in strains where only Mec1 is present; the observed signal likely reflects the small fraction of cells that have escaped α-factor arrest (Figures 1A and 1B).

**Figure 1:**
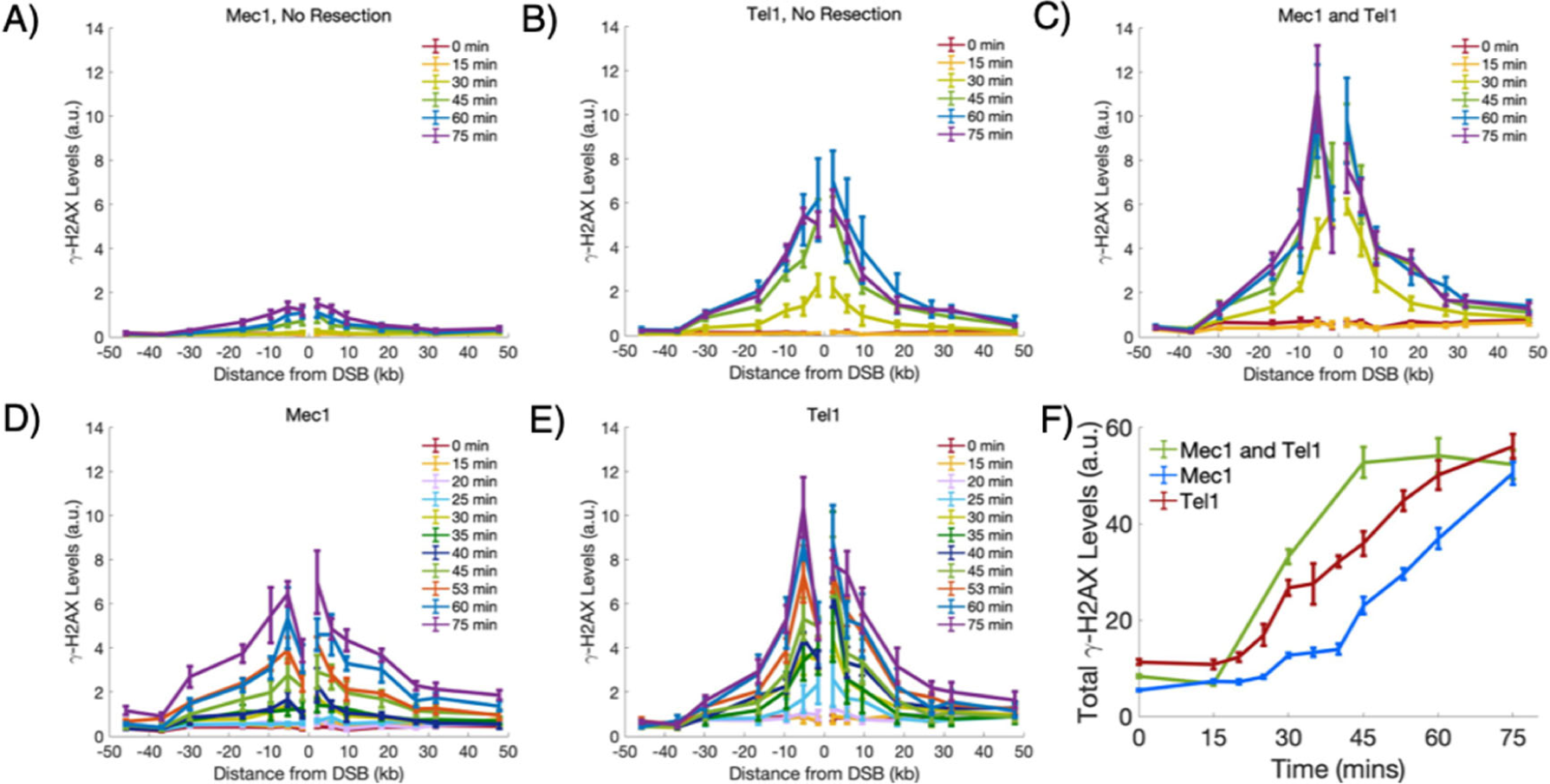
Experimental γ-H2AX profiles by Mec1 and Tel1. γ-H2AX spreading as measured by ChIP-qPCR after generating a break at the *MAT* locus. ChIP was performed using an antibody specific to γ-H2AX. All cells were arrested in G1 with end-joining prevented by *nej1*Δ or *yku80Δ*. In all plots, error bars represent standard error of the mean from n≥3 measurements. **A)** γ-H2AX spreading by Mec1 in *nej1*Δ cells. **B)** γ-H2AX spreading by Mec1 in *nej1*Δ cells. **C)** γ-H2AX spreading by both Mec1 and Tel1 in *ku80Δ* cells. **D)** γ-H2AX spreading by Mec1 in *ku80Δ* cells. **E)** γ-H2AX spreading by Tel1 in *ku80Δ* cells. **F)** Total γ-H2AX levels by both Mec1 and Tel1 (green), only Mec1 (blue) and only Tel1 (red) in *ku80Δ* cells.

To compare Mec1 and Tel1 activity in G1 cells, we deleted *YKU80*, which normally blocks access of the exonuclease Exo1 to the DSB (20). *yku80*Δ cells generate sufficient ssDNA to recruit the single-strand binding protein complex RPA and activate Mec1-Ddc2 (14). Under these conditions, both Mec1 alone or Tel1 alone were efficient at phosphorylating histone H2A (Figures 1D and 1E). In the absence of *YKU80*, the Tel1-only derivative exhibited a significant increase in γ-H2AX levels over background by 25 min (Figure 1E), when about 60% of the *MAT* locus had been cleaved (Figure S1). The appearance of γ-H2AX in the Mec1-only strain was slightly delayed by about 5 min (Figure 1D). In both cases, the extent of phosphorylation was largely complete by 75 min, but with noticeable differences between the phosphorylation profiles of Mec1 and Tel1. Although Tel1 appears to be more rapidly activated, the extent of γ-H2AX spreading for Tel1 is more confined to the region adjacent to the break than Mec1. The difference in the profiles of modification can be shown by calculating the mean modification distance (MMD), the distance from the break that encompasses half the γ-H2AX profile. At 75 min, the 95% confidence interval (CI) for the MMD of Mec1 is MMD_Mec1_= [12.9, 14.5] kb while the 95% CI for the MMD of Tel1 is MMD_Tel1_= [10.8, 11.6] kb, indicating that the activities of Mec1 and Tel1 lead to significantly different profiles of spreading (Figure S2A).

When both kinases are active, the kinetics of phosphorylation were more rapid and reached an apparent steady state around 45 min, suggesting that both Mec1 and Tel1 participate in phosphorylating H2A (Figures 1C and 1F). We also note that at later time points, the level of modification close to the DSB does not increase, most likely due to loss of nucleosomes during 5’ to 3’ resection that displaces histones, at a rate of 4 kb/hr (21). Although the γ-H2AX profiles for each kinase alone or when both are active differ in their kinetics and extent of spreading, the total amount of γ-H2AX formation is the same by 75 min, suggesting that Mec1 and Tel1 have similar levels of phosphorylation activity, but distribute the γ-H2AX sites differently (Figures 1F and S2B).

### The amount, but not the profile, of γ-H2AX is affected by reducing the density of phosphorylation sites

To further investigate the differences between the Mec1 and Tel1 modes of spreading, we mutated the phosphorylation site of one or the other H2A gene. Using CRISPR/Cas9, we mutated either *HTA1-S129* or *HTA2-S129* to alanine (see Methods), so that the density of phosphorylatable H2A-S129 sites should be reduced by half. As expected, each of these mutations reduced the level of γ-H2AX (Figures 2A, 2B and S2B) suggesting that the availability of H2A phosphorylation sites is limiting – i.e. the amount of γ-H2AX formed depends on the density of phosphorylatable sites. Despite the reduction in γ-H2AX levels, the extent of phosphorylation spreading for Mec1 and Tel1 remain unchanged when only 50% the sites can be phosphorylated; moreover, the two mutant profiles were not significantly different from each other. The 95% CI of the MMDs at 75 min for Mec1 are MMD_Mec1,HTA1_=[13.2, 14.4] kb and MMD_Mec1,HTA2_= [13.0, 14.2] kb, while the 95% CI of the MMDs for Tel1 are MMD_Tel1,HTA1_= [10.0, 10.8] kb and MMD_Tel1,HTA2_= [10.7, 11.9] kb (Figures 2 and S2A).

**Figure 2:**
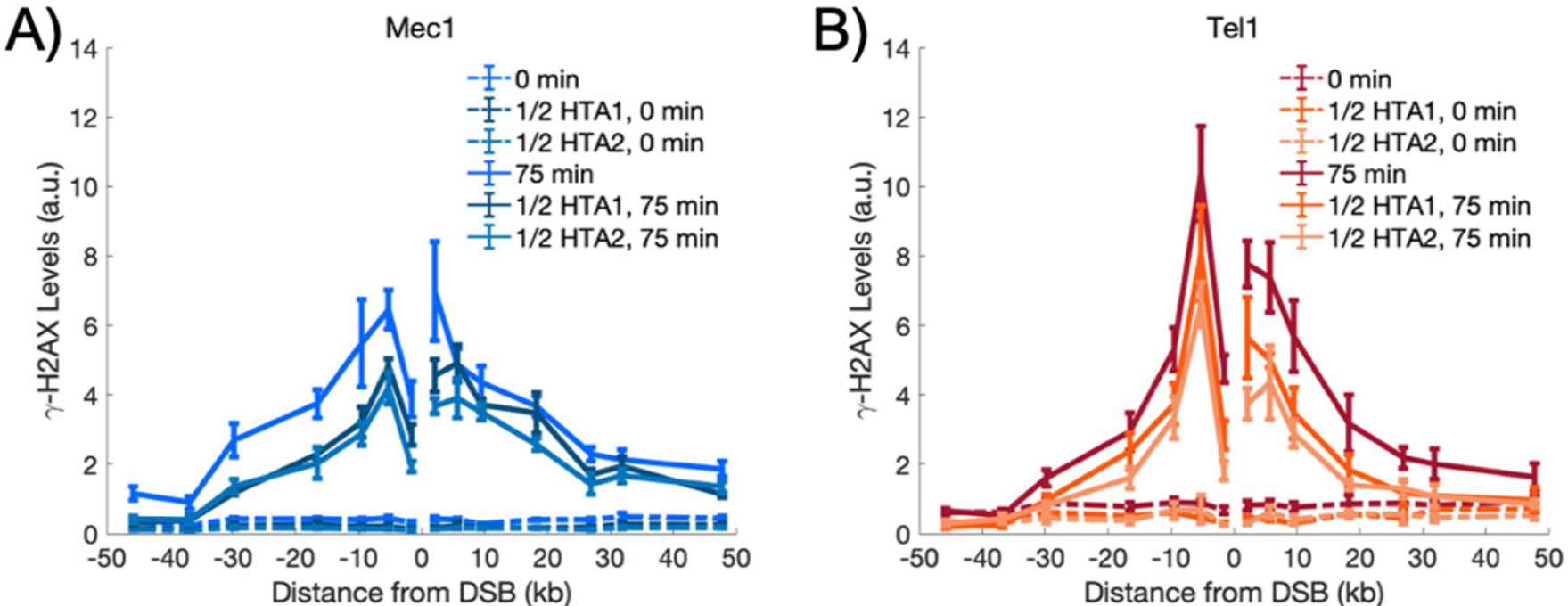
Experimental γ-H2AX profiles by Mec1 and Tel1 when 50% of H2A sites cannot be phosphorylated. Phosphorylation spreading in strains where one of the two *HTA* genes is mutated to *HTA-S129A*, rendering half the H2A sites non-phosphorylatable. Experimentally measured γ-H2AX profiles in the presence 50% phosphorylatable H2A by Mec1 **(A)** or Tel1 **(B)** are shown for 0 min (dashed lines) and 75 min (solid lines). “1/2 *HTA1*” refers to the presence of an *hta2-S129A* mutation, while “1/2 *HTA2*” carries the *hta1-S129A* mutation. Error bars represent standard error of the mean from n≥3 measurements.

### Mec1 modifies histones at distant chromosome sites that are recruited to the DSB

We previously showed that a DSB near one centromere will lead to the modification of all the other 15 pericentromeric regions clustered at the spindle pole body. This modification is predominantly carried out by Mec1 and occurs at about 1/10 the magnitude as modifications on the broken chromosome (22). γ-H2AX is also weakly spread around the recombination enhancer, RE, a sequence near the *HML* locus and roughly 170 kb from *MAT*, that facilitates pairing between *MAT***a** and the *HML* donor (23). RE binds multiple copies of the Fkh1 protein whose FHA domain can presumably also bind to phospho-threonines that are generated near the DSB; however, the specific phosphorylated target of the FHA domain remains unknown. ChIP using an anti-γ-H2AX antibody pulls down the RE region in *MAT***a** cells, when Fkh1 binds to RE, but not in *MAT*α cells, when Fkh1 binding is repressed (24). Thus, a kinase originating at the *MAT* locus can only phosphorylate the region around RE by 3D diffusive or looping mechanisms, and only when it has been brought into proximity with the DSB. In our strains *HML* is deleted, but the Fkh1 proteins bound to RE are still able to interact with phosphorylated targets near *MAT*. At 75 min, the level of ChIP-qPCR 10 kb away from RE was significantly increased above background for Mec1 (Figure 3A), while Tel1 did not show a significant increase at these distances, except possibly at the −5 kb location (Figure 3B). Taken all together the experimental evidence suggests that Mec1 is the kinase that is primarily responsible for phosphorylation spreading in *trans*.

**Figure 3:**
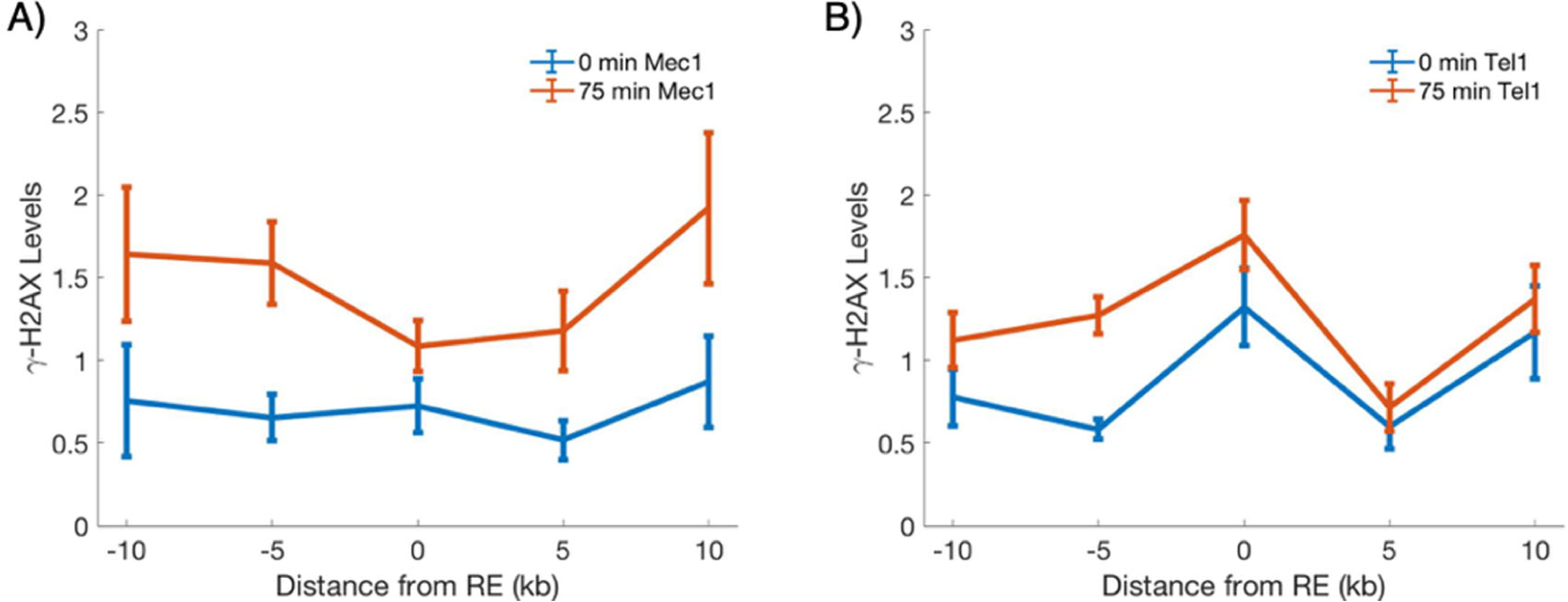
γ-H2AX measurements around RE. γ-H2AX levels measured around the recombination enhancer (RE), a locus on ChrIII known to interact with the MAT locus by forming a chromatin loop. **A)** γ-H2AX formation around RE by Mec1. Error bars represent standard error of the mean from n≥4 measurements. **B)** γ-H2AX formation around RE by Tel1. Error bars represent standard error of the mean from n≥4 measurements.

### Biophysical models of chromatin-modification spreading can be used to determine the mechanism by which Mec1 and Tel1 phosphorylate distant H2As

By comparing the phosphorylation data to mathematical models of phosphorylation spreading, we can infer the mechanisms by which the kinases spread γ-H2AX. We focused on two classes of models, one of which assumes that the spreading occurs by the kinase moving three-dimensionally through the nucleoplasm while the other assumes that the kinase moves one-dimensionally along the chromatin. For each class, we chose two minimal models commonly found in the literature (25): the 3D models are represented by chromatin looping and 3D diffusion while the 1D models are represented by 1D diffusion and directed sliding (Figures 4A-4D). (The models are described in detail in *Computational Methods* and *Supplementary Information*). All four models assume an initial recruitment of the kinases to the break site with a rate *k*_*init*_ but differ thereafter. For the 1D models, the parameter *k*_*init*_ comprises both the recruitment and detachment from the DSB to begin traversing the chromosome. Although each nucleosome contains two monomers of H2A, in these models we treat each phosphorylation site as a separate entity.

**Figure 4:**
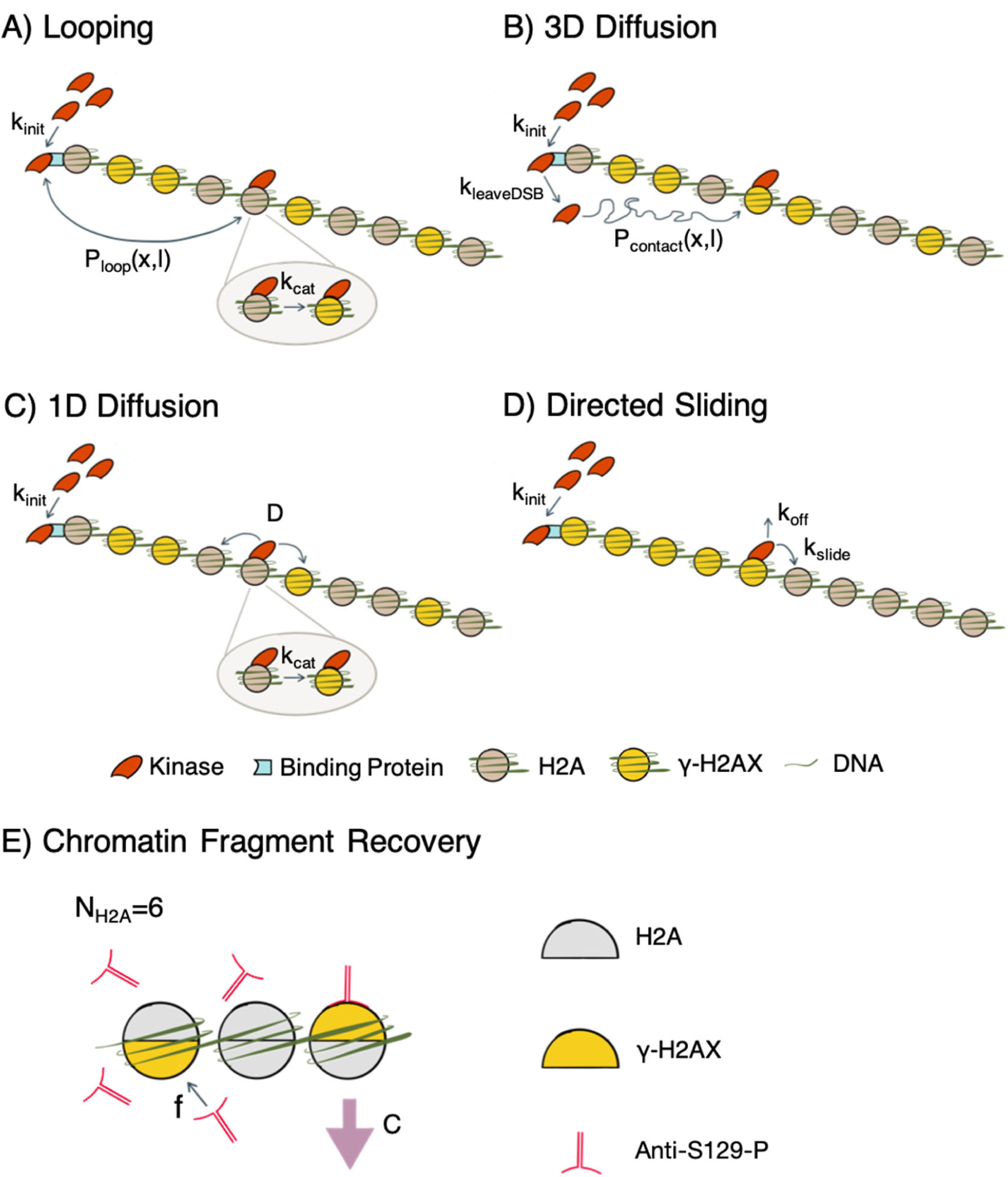
Schematic of phosphorylation spreading mechanisms. Schematics of looping **(A)**, 3D diffusion **(B)**, 1D diffusion **(C)** and directed sliding **(D)** mechanisms are depicted. The kinases Mec1 and Tel1 (red) are recruited to the DSB at a rate *k*_*init*_ to RPA and MRX respectively (teal). For the one dimensional models, 1D diffusion and directed sliding, *k*_*init*_ comprises both the kinase’s recruitment to and detachment from the DSB to begin translocating along the chromatin. The kinase proceeds to phosphorylate H2A (grey) to form γ-H2AX (yellow) using one of the four mechanisms. **A)** In the looping model, the kinase remains tethered at the DSB and forms a looped conformation with probability *P*_*loop*_(*x,l*), where *x* describes the distance of the target H2A from the break and *l* is the Kuhn length of the chromatin. *P*_*loop*_(*x,l*) is formulated using a worm-like chain polymer model. The kinase phosphorylates the H2A at a rate *k*_*cat*_ after contact is established with the H2A. ***B*)** In the 3D diffusion model, the kinase transiently binds to the DSB where it is activated and released at a rate *k*_*leaveDSB*_. The activated kinase diffuses away from the DSB until it encounters and phosphorylates an H2A. *P*_*contact*_(*x,l*) is the probability that the kinase will come into contact with an H2A located a distance x away from the break. ***C*)** In the 1D diffusion model, the kinase is recruited to the DSB and proceeds to diffuse along the chromatin with a one-dimensional diffusion coefficient *D*. The kinase phosphorylates the H2A that it comes into contact at a rate *k*_*cat*_. ***D*)** In the directed sliding model, the kinase moves unidirectionally away from the DSB along the chromatin. The kinase slides onto adjacent histones at a rate *k*_*slide*_ and phosphorylates all H2As encountered until the kinase detaches from the chromatin at a rate *k*_*off*_. ***E*)** Sonication during ChIP yields chromatin of ∼500bp. The parameter *N*_*H2A*_ describes the average number of H2As (grey) present on the DNA fragment. In this depiction, N_H2A_=6. Each γ-H2AX (yellow) is bound by an antibody (red) with probability *f*. Antibody binding to one γ-H2AX is independent of the other H2A sites on the same chromatin fragment. The binding of one antibody is sufficient for the pull down of the entire fragment (purple arrow). The recovery of the fragment is multiplied by the parameter *C* to account for the loss of DNA during the wash steps in ChIP.

Thermal fluctuations cause chromosomes to form dynamic conformations, including transient loops. In the looping model (Figure 4A), the kinase is tethered at the break end and folding of the chromatin brings distant H2As into physical contact with the DSB-bound kinase. Using a worm-like chain model of the chromosome, we compute *P*_*loop*_(*x,l*), the probability of a looped conformation in which the DSB-bound kinase is in contact with an H2A located *x* kilobases away, where *l* is the Kuhn length of the chromatin. When the kinase is in contact with an H2A, phosphorylation occurs at a rate *k*_*cat*_.

The 3D diffusion model (Figure 4B) assumes that the kinase transiently binds to the DSB, where it is activated, and then released at a rate *k*_*leaveDSB*_. The activated kinase diffuses through the nucleoplasm until it encounters and phosphorylates an H2A site. For an H2A located *x* kilobases from the DSB, we use the worm-like chain polymer model to calculate the mean 3D distance between this H2A and the DSB. This distance is used to compute the probability *P*_*contact*_(*x,l*) that the kinase will come into contact with the H2A.

In the 1D diffusion model (Figure 4C), we assume that the kinase lands at the DSB and proceeds to diffuse along the chromatin with the one-dimensional diffusion coefficient *D*. When it encounters a histone, the kinase phosphorylates the H2A at a rate *k*_*cat*_. The directed sliding model (Figure 4D) is similar to 1D diffusion except that the kinase moves unidirectionally away from the DSB along the chromatin. The kinase slides onto adjacent histones at a rate *k*_*slide*_ and phosphorylates all H2As that it comes across until the kinase detaches from the chromatin at a rate *k*_*off*_.

For an H2A located *x* kilobases from the break site, each of our models predicts the probability *P*(*x,t*) that the histone has been phosphorylated at time *t* after the DSB induction. Each model predicts a distinct *P*(*x,t*) allowing us to use the experimentally measured γ-H2AX profiles to determine the best spreading models for Mec1 and Tel1. However, before we can directly compare the theoretical predictions to the experimental measurements, it is necessary to convert the predicted probabilities *P*(*x,t*) to the expected ChIP signals. We formulate a simple thermodynamic model for the ChIP pulldown to account for the binding of antibodies to a chromatin fragment containing multiple phosphorylated H2As (Figure 4E). Sonication during ChIP results in chromatin fragments of roughly 500 bp. We introduce the following model parameters: *N*_*H2A*_, *f* and *C*. The parameter *N*_*H2A*_ represents the average number of H2As on the chromatin fragment, and *f* is the probability that each γ-H2AX is bound by an antibody; we assume that the γ-H2AX-antibody interaction is independent of the other γ-H2AXs on the same chromatin fragment. We also make the assumption that the presence of one γ-H2AX-antibody interaction is sufficient for pulling down the entire chromatin fragment during ChIP. Finally, *C* accounts for the loss in DNA recovery during the wash steps of ChIP. We derive the ChIP model in detail in the *Supplementary Information*.

### Bayes factors reveal the most likely phosphorylation spreading mechanisms for each kinase

The best model for each kinase was determined by calculating the Bayes factor, which expresses how much less probable one model is compared to another model, given all the data collected in our experiments. When evaluating the Bayes factor, we take into account both the γ-H2AX profiles around the DSB (Figures 1D and 1E) as well as the phosphorylation in *trans* near the RE locus (Figure 3). The results are recorded in Table 1 as the log_10_ of the Bayes factor. Using Bayesian model selection, we determined that 3D diffusion is the most likely mechanism for Mec1, while directed sliding is the most likely mechanism for Tel1.

**Table 1:** log_10_(Bayes Factor) for various model combinations. log_10_(Bayes Factor) is shown for each model. ***A*)** Bayes factors were computed for the four mechanisms from Tel1-mediated phosphorylation. 1D mechanisms are much more likely than 3D mechanisms for Tel1, with directed sliding as the most likely mechanism. The Bayes factor was calculated by dividing the probability of the indicated model by the probability of the best model. For instance, −5 in the second row means that the probability of Tel1 undergoing 1D diffusion is 10^−5^ times less likely than directed sliding. ***B*)** Bayes factors were computed for Mec1 and Tel1 simultaneously. The most likely model combination is Mec1 3D diffusion and Tel1 directed sliding.

| Model for Tel1 | $\log_{10}$(Bayes Factor) |
| --- | --- |
| Directed Sliding | Best Model |
| 1D Diffusion | -5 |
| 3D Diffusion | -61 |
| Looping | -251 |

**B)**
| Model Combination | | $\log_{10}$(Bayes Factor) |
| --- | --- | --- |
| Mec1 – 3D Diffusion; | Tel1 – Directed Sliding | Best Model |
| Mec1 – 3D Diffusion; | Tel1 – 1D Diffusion | -6 |
| Mec1 – 1D Diffusion; | Tel1 – Directed Sliding | -12 |
| Mec1 – 1D Diffusion; | Tel1 – 1D Diffusion | -18 |
| Mec1 – Directed Sliding; | Tel1 – 1D Diffusion | -20 |
| Mec1 – Directed Sliding; | Tel1 – Directed Sliding | -35 |
| Mec1 – Looping; | Tel1 – Directed Sliding | -259 |
| Mec1 – Looping; | Tel1 – 1D Diffusion | -258 |

We first computed the Bayes factors for Tel1 alone (Table 1A). It is clear that 1D models are vastly more likely than the 3D models for Te1l. Therefore, we discarded the 3D models for Tel1. Next, we simultaneously computed the Bayes factors for both 1D models of Tel1 and either 1D or 3D models for Mec1 by imposing the constraint that the values of the ChIP-model parameters (*C, f* and *N*_*H2A*_) must be the same regardless of which kinase is active. The simultaneous pairwise calculations of the Bayes factor for both Mec1 and Tel1 models (excluding the 3D models for Tel1) are shown in Table 1B. Further details of this calculation can be found in the *Supplementary Information*.

After determining the best models for Tel1 and Mec1, we implemented Bayesian parameter estimation to find the optimal values for the model parameters. When performing parameter estimation, the data in Figures 1D, 1E, 2A, 2B, 3A, and 3B were fit simultaneously to the best models. The best models are plotted using the optimal parameter values and are in quantitative agreement with the experimental data (Figure 5). All of the less likely models are plotted against data in Figures S3-S14.

**Figure 5:**
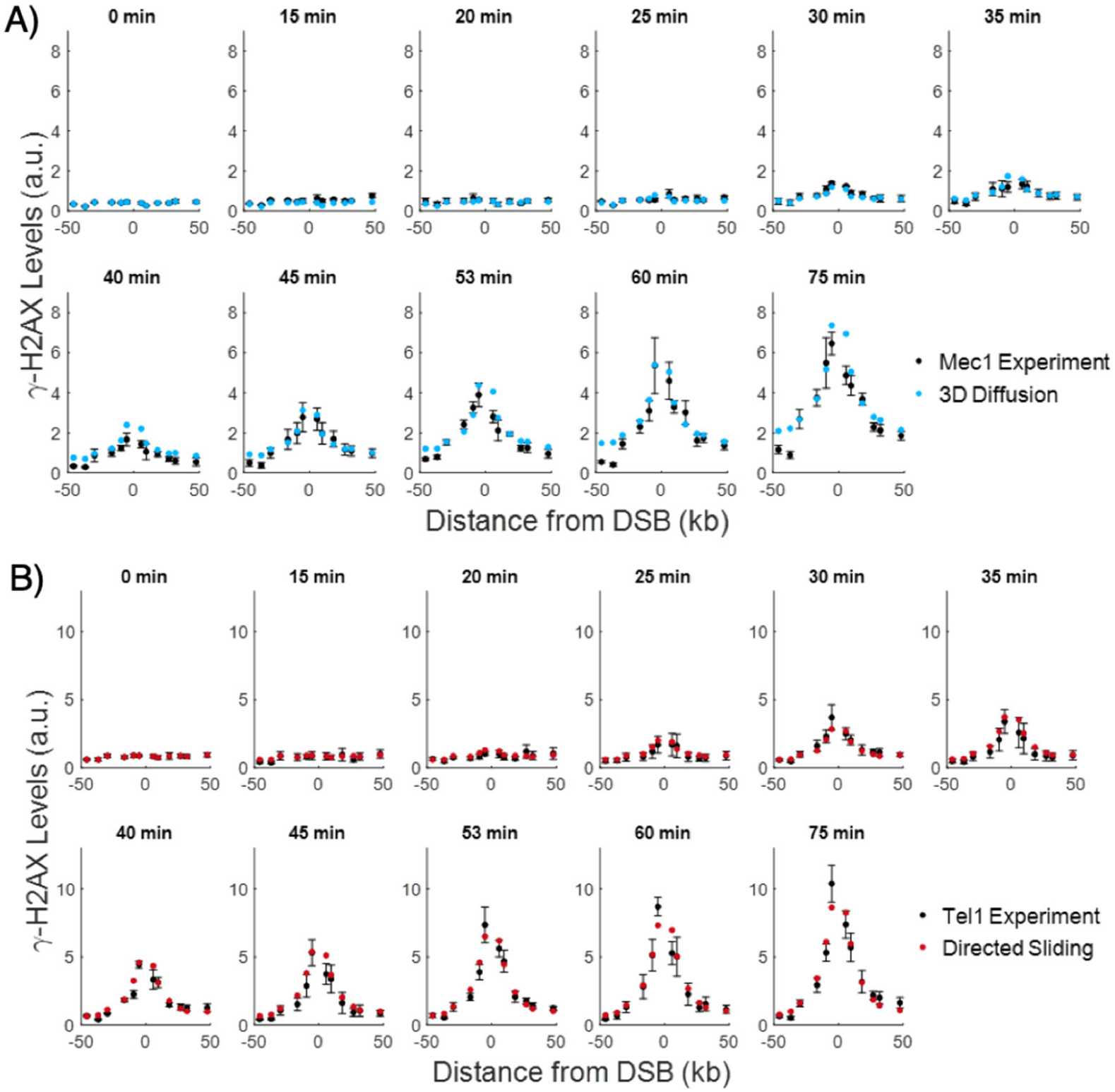
Comparison of experimental γ-H2AX profiles to the most likely theoretical spreading distributions. Comparison of experimental γ-H2AX profiles (black) to theoretical γ-H2AX distributions for the best model. Experimental and theoretical curves for 3D diffusion (blue) by Mec1 phosphorylation are shown in (**A)** and experimental and theoretical curves for directed sliding (red) by Tel1 are shown in (**B)**. The experimental error bars represent the standard error of the mean from n≥3 measurements. The theory curves are plotted using parameters values shown in Table 2. Due to concerns about the reduction in γ-H2AX signal from resection close to the DSB, the −1.6 kb and −2.1 kb data points (not shown) were excluded when performing fits to the experimental data. For the plots in **A** and **B**, our theoretical predictions start from the same background levels as in the experimental data.

In Figure 6A, we plot the predicted phosphorylation levels by Mec1 around RE. The 3D models predict that when RE is bound to *MAT*, the loci within 10 kb of RE will be in close proximity to the DSB and so the kinase will phosphorylate H2As at these loci. When modeling γ-H2AX spreading around RE, we include two additional parameters, which take into account the binding of RE to *MAT* (see *Supplementary Information*). We did not plot the predicted phosphorylation by Tel1 near RE because the directed sliding model predicts that RE is too far away from the DSB for there to be any phosphorylation by Tel1 above the background level.

**Table 2:** Parameter values for the best models. Displayed are the optimal parameter values obtained when Bayesian parameter estimation was performed simultaneously for Mec1 and Tel1. The 95% CI for the parameter values are shown in parentheses. A description of the parameters can be found in *Computational Methods*.

| Tel1, Directed Sliding | Mec1, 3D Diffusion | Parameters Shared by<br>Mec1 and Tel1 |
| --- | --- | --- |
| $k_{init}$: 0.033/minute (0.0060-0.077) | $k_{init}$: 0.089/minute (0.068-0.15) | $C$: 26 (13-280) |
| $k_{slide}$: 3.2 kb/minute (2.0-17) | $l$: 14 kb (9.9-15) | $f$: 0.16 (0.020-0.72) |
| $k_{off}$: 0.30/minute (0.17-1.8) | $\omega$: $4.2 \times 10^{-3}$/minute<br>($1.1 \times 10^{-3}$-$5.0 \times 10^{-3}$) | $N_{H2A}$: 6 (2-6) |

**Figure 6:**
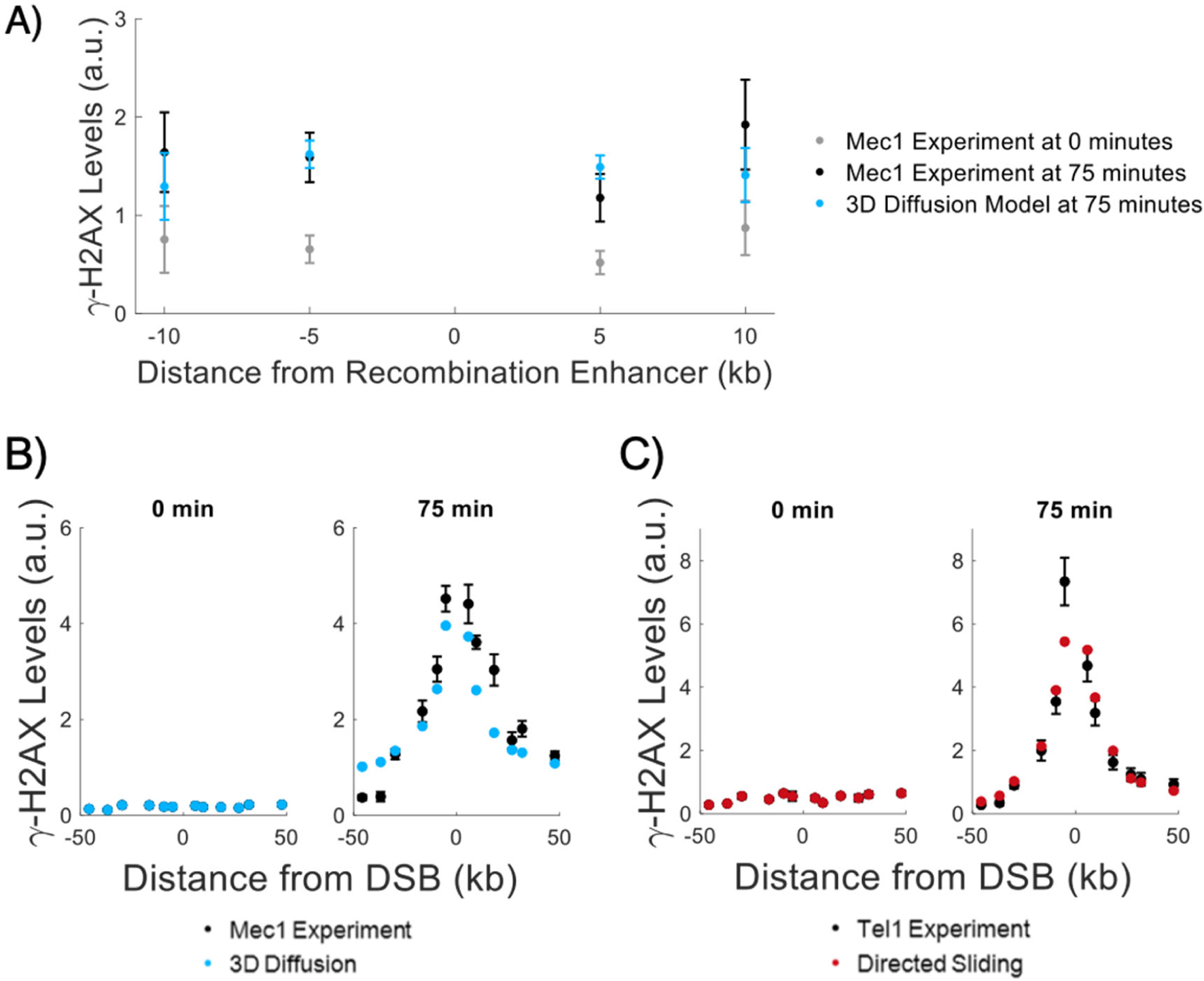
Comparison of experimental γ-H2AX profiles to theoretical predictions around RE and in strains in which 50% of H2A sites cannot be phosphorylated. Comparison of the 3D diffusion model (blue) with the γ-H2AX measurements around the Recombination Enhancer (RE) by Mec1 is shown in (**A)**. The experimental data at 0 min (grey) and 75 minutes (black) are displayed along with theoretical predictions based on the 3D diffusion model (blue). Experimental error bars are the standard error of the mean. Theoretical predictions from 3D diffusion were generated using the parameters listed in Table 2 with the exception of *k*_*init*_ = 0.025/minute, to account for an initial delay is RE binding, consistent with the kinetics of mating type switching as measured in Avşaroğlu et al. (24). Also, the RE-DSB interaction is assumed to occur only 19% of the time. The data point at RE was excluded from the plot since the theory does not make predictions at this point. The model starts from the same background levels as the experimental data, so the large signal to noise ratio at 0 min leads to large error bars at 75 min in the theoretical curve. The experimentally measured γ-H2AX distributions for the *HTA-S129A* mutants were averaged together for each kinase. In (**B)** and (**C)**, theoretical curves were generated using the parameters listed in Table 2 for 3D diffusion by Mec1 (blue) and directed sliding by Tel1 (red), respectively, and overlaid onto the experimentally measured 50% γ-H2AX profile (black).

### Theoretical predictions of phosphorylation spreading are in agreement with data from mutant strains in which 50% of the H2As are phosphorylatable

Next, we compared the measured γ-H2AX levels from the 50% phosphorylatable strains to the theoretical predictions of the 3D diffusion model for Mec1 and the directed sliding model for Tel1 (Figures 2, 6B and 6C), when only one out of every two H2As can be phosphorylated. For the 50% phosphorylatable strains, we used the same parameter values as those used to fit the wild-type H2A strains (Figure 5). The theoretical predictions for the 50% phosphorylatable strains are largely in agreement with the experimental data (Figures 6B and 6C).

While it was conceivable that the introduction of the S129A mutant allele might impact the sliding rate of Tel1, we find that, in fact, Tel1’s sliding remains unaffected. We performed fits in which the *HTA-S129A* strains were allowed to have both *k*_*slide*_ and *k*_*off*_ differ from the parameter values established for the wildtype H2A strains. The sliding parameters remain unchanged in the *HTA-S129A* mutants with the optimal *k*_*slide*_ and *k*_*off*_ values being within 10% of the established rates for wildtype H2A strains. Moreover, the Bayes factor calculation tells us that the most likely directed sliding model for Tel1 is one in which *k*_*slide*_ and *k*_*off*_ are unaltered in the 50% phosphorylatable strains. From our Bayesian parameter estimation, the average distance over which Tel1 slides is 11 kb (95% CI of 9.7–12 kb) before falling off the chromosome.

The agreement between theory and experiments on *HTA-S129A* mutants also validates our model of the ChIP process, which provides us with a quantitative understanding of the relationship between the amount of γ-H2AX on the chromatin and the DNA recovery from ChIP (Figure 4E). The recovery of > 50% of the ChIP signal when one H2A gene is rendered nonphosphorylatable (Figure S2B) is explained by the assumption that there is a substantial chance that a DNA fragment will be pulled down even if there is only one γ-H2AX on the ∼500 bp DNA fragment (Figure 4E). Even though the *H2A-S129A* strains have half as many γ-H2AX per fragment, there is still enough γ-H2AXs per fragment to be recovered with ChIP (see *Supplementary Information*).

### Phosphorylation levels by Mec1 is improved by overexpressing its binding partner Ddc2

Previously Ddc2 overexpression was shown to increase Mec1’s checkpoint activity, whereas increasing Mec1 expression had no consequence (26). We therefore asked if the efficiency of H2A modification by Mec1 could be improved by increasing the concentration of Ddc2. We integrated plasmid PML105.45, which contains an extra copy of Ddc2 under a *GAL1* promoter, at *leu2-3,112* in the *tel1*Δ *yku80*Δ strain (26). Indeed, increased expression of Ddc2 in this Mec1-only strain increased the amount of γ-H2AX (Figure 7A). In this instance, the total amount of γ-H2AX by Mec1 at 60 min in the Ddc2 overexpressed strains is nearly the same as that measured for the Tel1-only strain (Figure 7B). However, the distribution of γ-H2AX sites is unaltered by the overexpression of Ddc2 since the MMD was unchanged from that seen in the absence of Ddc2 overexpression (Figure S2A). The 95% CI for the MMD for the Ddc2 overexpression case is given by MMD_Mec1,Ddc2 O/E_= [12.5, 14.1] kb while the MMD for Mec1 is MMD_Mec1_= [12.9, 14.5] kb.

**Figure 7:**
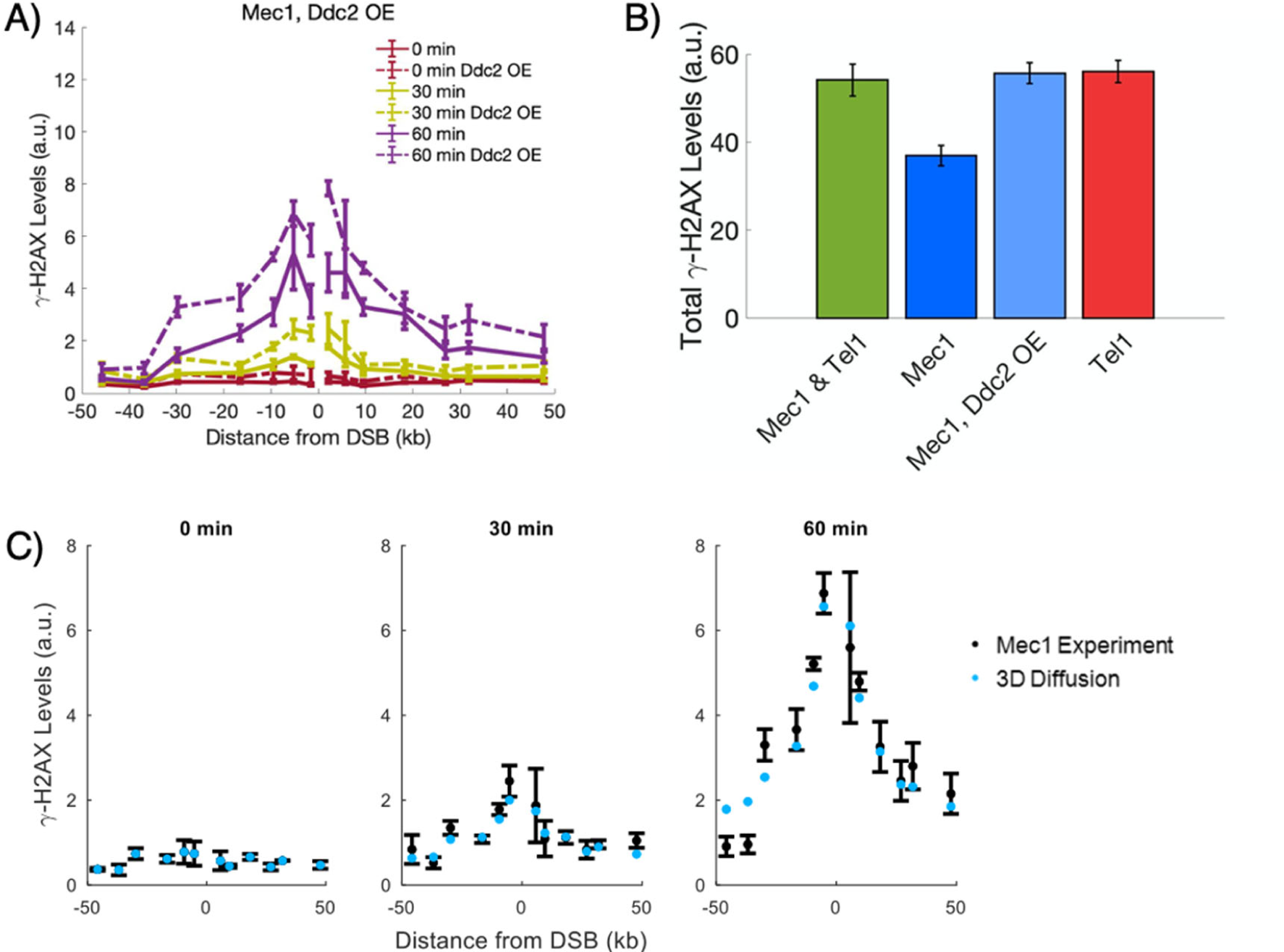
γ-H2AX profiles by Mec1 with Ddc2 overexpression. **A)** Overexpression of Ddc2 leads to increased γ-H2AX levels by Mec1. Experimental error bars represent the standard error of the mean from n=2 measurements. **B)** The overexpression of Ddc2 increases the total amount of phosphorylation by Mec1 (blue compared to light blue). By 60 min, Ddc2 overexpression rises the amount of γ-H2AX formed to those of Tel1-mediated phosphorylation levels (light blue compared to red). Error bars were calculated by error propagating over the error of the γ-H2AX profiles at 60 min for the associated strain. **C)** The prediction from the 3D diffusion model (blue) agrees with the experimental γ-H2AX levels (black) for Mec1-mediated phosphorylation in the presence of Ddc2 overexpression. Experimental error bars represent the standard error of the mean from n=2 measurements. Predicted levels of γ-H2AX were calculated by treating the initial recruitment rate *k*_*init*_ as a free parameter and by using the optimal parameters found for Mec1-mediated phosphorylation under wildtype levels of Ddc2 expression (Table 2). By performing Bayesian parameter estimation, we find that *k*_*init*_ in the Ddc2 overexpressed case is 2.8 times higher than the optimal value in the wildtype Ddc2 case.

To determine the effects of Ddc2 overexpression and further test the 3D diffusion model of phosphorylation spreading by Mec1, we compared the theoretical model to the Ddc2-overexpressed strain. Since Ddc2 forms an obligate heterodimer with Mec1 prior to its loading onto RPA-coated ssDNA (23, 24), the overexpression of Ddc2 should lead to an increased rate of Mec1 recruitment, *k*_*init*_. Treating *k*_*init*_ as a free parameter, while constraining all other model parameters to their previous values, we find that the 3D diffusion model agrees with the experimental data from the Mec1 strains in which Ddc2 is overexpressed (Figure 7C). The optimal *k*_*init*_ is 0.25/minute, 2.8 times the optimal *k*_*init*_ for the Mec1-only strains where Ddc2 is expressed at wild-type levels.

## Discussion

Although we know a great deal about the evolutionarily related phosphoinositol-3-kinase like kinases ATM and ATR, including their structure and phosphorylation targets (29), the manner by which these kinases reach their targets have not been well characterized. In this paper, we used Bayesian model selection to distinguish how Mec1 and Tel1 are able to create extended regions of γ-H2AX after DNA damage. Our analysis shows that the experimental profile for Mec1 is best matched by a 3D diffusive mechanism. However, we note that the 1D models also produce good fits to the Mec1 γ-H2AX profile around the DSB (Figures S6-S8). Therefore, we do not rule out the possibility that Mec1 undergoes some 1D diffusion along the chromatin in addition to 3D diffusion. Meanwhile, the narrow shape of Tel1’s γ-H2AX profile around the DSB is best fit by a directed sliding model and poorly fit by the 3D-spreading models (Figures 5B, S9-S14).

The γ-H2AX profiles also provide us with insights into the recruitment process of the kinases. Since Tel1 is recruited to the nearly blunt ends created by HO endonuclease (13), whereas Mec1 requires that there be some 5’ to 3’ resection of the end (30), Tel1 would be expected to be activated before Mec1 and indeed this appears to be the case (Figures 1D-1F). However, when we increased the expression of Ddc2, the kinetics of Tel1 and Mec1 activation were nearly identical, as the total γ-H2AX levels are now comparable by 60 min (Figure 7B). As Ddc2 is necessary for Mec1 loading onto RPA near the DSB, this result suggests that it is Mec1 binding to RPA rather than a delay in resection that limits the initial response.

One might also expect that, due to Tel1’s association with the MRX complex, Tel1 levels should increase in the absence of resection since MRX is recruited to nearly blunt DNA ends, but this was not found to be the case (Figures 1B and 1E). We found that the amount of γ-H2AX formation was somewhat reduced in G1-arrested *nej1*Δ cells (where resection is impaired) compared to those in *yku80*Δ cells (Figure S2B). It should be noted that there must be a number of MRX complexes recruited to the DSB in order that GFP-tagged proteins can be seen as a robust focus (31). It is possible that MRX-Tel1 binding is reduced by competitive binding of yKu70-yKu80, which also preferentially binds to dsDNA ends (32).

Previous studies have performed ChIP measurements of Mec1-Ddc2 and Tel1 adjacent to the DSB, finding that they increase roughly linearly over a time span longer than our experiments (33, 34). While these results could indicate an increasing number of kinase copies at the break site, our Bayes factor calculations suggest that that even with increasing kinase occupancy, the total kinase activity saturates – i.e. the rate of phosphorylation is constant over the 75 min of our experiment, not increasing. This was determined by considering both a constant phosphorylation rate and an increasing phosphorylation rate for all spreading models; we found that the γ-H2AX profiles were best fit when the rate of phosphorylation remained constant over time (see Table S1 and *Supplementary Information, Model Derivations*). The constant rate is assumed for all models in the main text.

The initial recruitment of Mec1 to ssDNA only confines it to sites within a few kb of the DSB end and does not account for the rapid spreading of γ-H2AX down the chromosome, since the rate of resection of DSB ends is only 4 kb/hr (21). We note that while γ-H2AX spreads rapidly over 50 kb from the break within an hour, H2A modifications continue to spread much more slowly over at least another 50 kb, so that modifications around *CEN3* – 100 kb from the DSB at *MAT* – are seen only after 8-10 h (35). This slower rate of modification parallels the rate of 5’ to 3’ resection and proves to be performed only by Mec1 (36). This activity suggests that, as new ssDNA is generated by the inexorable action of 5’ to 3’ exonucleases, Mec1 might re-load at newly synthesized ssDNA/RPA and proceed to diffuse from the new ssDNA to extend γ-H2AX another 50 kb from the new site of entry.

Ddc2 assembles at the DSB in a Mec1-dependent manner (26, 27). Structural work on the Mec1-Ddc2 heterodimer suggests that it forms a stable focus at RPA-coated ssDNA with the coiled-coil domain of Ddc2 acting as a flexible linker so that Mec1 can freely phosphorylate nearby targets (28). If Mec1-Ddc2 remains at the break, then the looping mechanism would be the only possible option among the models that we considered, since the other mechanisms would require the kinase to detach from the DSB after recruitment. However, our analysis shows the predicted γ-H2AX profile from a looping mechanism does not fit the experimentally measured profile (Figures S3 and S4). In formulating the 3D mechanisms, we took into account the Kuhn length of the chromatin, as determined by both HiC and florescence microcopy experiments, and thus restricted the Kuhn length to 8.4-15 kb, the 95% CI for the Kuhn length reported by Arbona et al. (37). Under these constraints, 3D diffusion is the preferred Mec1 mechanism, while looping interactions are unfavorable. Moreover, a 3D mechanism is supported by γ-H2AX measurements around RE, accounting for Mec1’s ability to act in *trans* by phosphorylating histones on undamaged chromatin brought into close proximity with the DSB (Figures 3A and 6A). Additionally, Mec1 has been shown to phosphorylate histones clustered at pericentromeric regions when a DSB is generated close to one centromere (22).

Our conclusion of Tel1 sliding is seemingly at odds with prior results that support the idea that Tel1 is tethered to the break by the MRX complex, just as mammalian ATM is associated with MRN (38). ChIP measurements show that multiple copies of MRX (and presumably Tel1) remain associated with the DSB for 2-3 h (31), much longer than the 75 min duration of our experiments. However, our analysis has shown that looping, which we associate with a DSB bound kinase, is highly unlikely to be the mechanism employed by Tel1 since the theoretical predictions do not match the experimentally measured distribution and Bayesian analysis shows that 3D mechanisms are astronomically less probable than 1D sliding (Tables 1 and S1). Thus, although Tel1 is recruited to the DSB end, its subsequent action may not depend on the MRX complex.

The same mathematical formulation used to encode the directed sliding model can also be used to describe two other 1D spreading mechanisms: recruitment/assembly and loop extrusion. During recruitment/assembly, the arrival of a protein to the break facilitates the recruitment of subsequent proteins until a string of proteins is formed spanning many nucleosomes. In this scenario, the rate of recruitment of the next protein copy is analogous to *k*_*slide*_. This recruitment/assembly process has been suggested to occur in mammalian cells where the propagation of ATM down the chromosome is in part facilitated by the γ-H2AX binding proteins MDC1 and possibly 53BP1 (39–41). MDC1 is capable of recruiting the MRN complex, which in turn recruits ATM further from the break, which in turn forms γ-H2AX which act as MDC1 binding sites. The sequential assembly of MDC1 and ATM results in the propagation of γ-H2AX away from the break (33, 35). While yeast lacks a MDC1 homolog, it is possible that its 53BP1 homolog, Rad9, may play a similar role in Tel1 mediated γ-H2AX spreading (43).

Results similar to 1D sliding would also be predicted if chromatin near a DSB were actively extruded into a loop, a process different from the looping model in which the chromosome only undergoes passive looping due to thermal fluctuations. *In vitro* experiments have shown that chromatin can be actively pulled through the ring-shaped cohesin or condensin complexes, resulting in extruded chromatin loops that grow over time (29, 30). Indeed, in yeast, cohesin and another SMC complex, Smc5,6 are recruited to sites of DSB damage (46–48). An analogous role for condensin has not been examined. We imagine a mechanism in which Tel1 remains tethered to the break while cohesin (or another SMC complex) sits near the break site and pulls in nearby chromatin through the ring. The histones that are pulled through the ring would therefore slide past Tel1 and be phosphorylated. Hence, the rate of loop extrusion is analogous to the rate *k*_*slide*_ in the sliding model. From Bayesian parameter estimation for the sliding model for Tel1, we observe that the 95% CI for *k*_*slide*_ overlaps with the estimated range for the speed of loop extrusion for cohesin (44). Also, since the rate of loop extrusion is a property of cohesin, we expect that *k*_*slide*_ would be unaffected by the density of phosphorylatable sites in H2A. Consistent with this notion, we found that *k*_*slide*_ is unchanged in the strain where only 50% of the H2As can be modified. Furthermore, unlike the sliding mechanism, loop extrusion has the added benefit of tethering Tel1 at the break end resulting in a stable focus. On the other hand, it should be noted that our measurements were made with G1-arrested cells, where cohesin or other SMC complexes may not be able to assemble (49–51). We further note that the profile of γ-H2AX spreading we have observed here is not apparently different from our previous observations when a DSB was induced in logarithmically growing cells (12, 22).

To our knowledge these are the first detailed comparisons of the mechanisms of spreading of ATM and ATR phosphorylation of histone H2A. They reveal that these two proteins, despite their evolutionary relationship, have adopted different strategies to propagate a signal away from a broken chromosome end.

## Methods and Materials

### Strain Construction

Standard yeast genome manipulation techniques were used to construct all strains and linear DNA and plasmids were introduced using the standard lithium acetate transformation protocol (50, 51). All strains used were variants of the strain JKM139 (54). The yeast strains used in this study can be found in Table S2. All primer sequences and plasmids used during strain construction are listed in Tables S3 and S4.

*nej1*Δ::HPH mutants were constructed by amplifying pAG32 (HPH) using primers Nej1-MXp1 and Nej1-MXp2 and *yku80*Δ::HPH mutants were constructed by PCR-amplifying pJH1515 using primers ku80MX18 and ku80MX19 to create a fragment containing hygromycin-resistance with homology to *NEJ1* and *YKU80*, respectively. Linear DNA was introduced to the appropriate strain using lithium acetate transformation. *hta1-S129A* mutants were generated by transforming strains with pBL13, which contains *Cas9* and a gRNA targeting *HTA1*, and the repair template BL327. Similarly, *hta2-S129A* mutants were generated by transforming strains with pBL14, which contains *Cas9* and a gRNA targeting *HTA2*, and the repair template BL331. The exact primer sequences of the repair oligos are shown in Table S3. S129A mutations were confirmed through sequencing by GENEWIZ.

Ddc2 was overexpressed in strain yKL019 by integrating PML105.45 (obtained from Maria Pia Longhese) which carries a copy of *GAL1-Ddc2*. PML105.45 was cut with *Cla*I and integrated at *leu2-3,112* by transformation.

### Growth conditions and DSB induction

Cells from a single colony were grown overnight in 5 ml of YEPD, washed three times with YEP + 3% lactic acid (YEP-Lac) and then grown in 350 ml of YEP-Lac until log phase growth with a cell concentration between 5×10^6^ cells/ml and 8×10^6^ cells/ml. α-factor (United Biochemical) was added to the culture to a concentration of ∼5 µM and maintained in the cell culture for at least for two doubling times before cell collection. G1 arrest was confirmed microscopically. After G1 arrest, 20% galactose was added to the YEP-Lac culture to a final concentration of 2% to induce *GAL*::*HO* expression, resulting in cutting at the *MAT***a** locus.

### Chromatin immunoprecipitation

Cells were harvested from log-phase population before DSB induction and at various times after DSB induction. ChIP was carried out according to the protocol of Shroff et al. (9). Briefly, 45 ml of culture were fixed and crosslinked with 1% formaldehyde for 10 min, after which 2.5 ml of 2.5 M glycine was added for 5 min to quench the reaction. Cells were pelleted and washed 3 times with 4°C TBS. Yeast cell walls were disrupted by beating the cells with 425-600 μm glass beads for 1 h in lysis buffer at 4°C. The lysate was sonicated for 2 min to obtain chromatin fragments of ∼500 bp in length. Debris was then pelleted and discarded, and equal volume of lysate was immunoprecipitated using γ-H2AX antibody (abcam ab15083) for 1 h at 4°C, followed by addition of Protein-A agarose beads (Sigma-Aldrich #1719408001) for 1 h at 4°C. The immunoprecipitate was then washed twice in 140 mM NaCl lysis buffer, once with 0.5 M NaCl lysis buffer, once with 0.25 M LiCl wash buffer and once with TE. Crosslinking was reversed at 65°C overnight followed by proteinase-K and glycogen addition for 2 h. Protein and nucleic acids were separated by phenol extraction. LiCl was added to a final concentration of 400 mM LiCl. DNA was precipitated using 99.5% EtOH. A second precipitation step was carried out using 75% EtOH and the DNA resuspended in TE.

### Quantitative PCR

γ-H2AX levels around the DSB were assessed by quantitative PCR (qPCR) using the primer sequences listed in Table S5. qPCR of immunoprecipitated samples was carried out using a Rotor-Gene SYBR Green PCR Kit (Qiagen 204076) in a Qiagen Rotor-Gene Q real-time PCR machine. γ-H2AX levels at all distances around the break were normalized to input as measured at the *PHO5* locus with primers MT101 and MT102.

### Computational Methods

The predicted γ-H2AX profiles were simulated using the Gillespie algorithm (for the 1D diffusion model) or derived using master equations (for all other models). The models are described below and are derived in the *Supplementary Information*. There are 3 parameters specific to each model, plus an additional 3 parameters shared by all models. The shared parameters model the ChIP process.

All models begin with the formation of the DSB. Although induction of HO endonuclease is reasonably synchronous, the DSB does not form simultaneously in all cells, and we incorporate this variability into all models. As shown in Figure S1, the earliest cells incur a DSB was ∼12 minutes after galactose exposure, presumably the time required for the transcription and translation of HO endonuclease. Over time, the DSB formed in more cells and by 30 minutes, ∼80% cells exhibited a DSB. The percent of cells without a break follows an exponential decay (Figure S1). After DSB formation, all models include the arrival of kinases (Mec1 or Tel1) to the break site, occurring at a constant rate *k*_*init*_. For the 1D models, the parameter *k*_*init*_ comprises both the recruitment and detachment from the DSB to begin traversing the chromosome.

The 3D diffusion, 1D diffusion and directed sliding models require that the kinase becomes activated upon arrival to the break site. Without this activation, these models would not predict preferential phosphorylation of H2As close to the break site (see the *Supplementary Information* for exceptions to this requirement and a further discussion of activation). A precedent for such activation can be found in an *in vivo* study of ATM, the mammalian homolog of Tel1, in which ATM autophosphorylation and activation occurs through its association with the MRN complex (55). In budding yeast, Mec1 phosphorylates sites on its partner, Ddc2, but only does so after DSB formation, presumably when it is bound to the break region (14). For simplicity, the initial recruitment rate parameter *k*_*init*_ also encompasses the activation step.

#### Looping Model

We model the chromosome as a worm-like chain at thermodynamic equilibrium, which has previously been shown to predict the frequencies of physical contact between chromosomal loci in yeast (56) (and similar polymer models have accurately predicted contact frequencies as well as the 3D positioning of genes in the yeast nucleus (24, 37, 57)). The thermodynamic equilibrium assumption is justified because we estimate that loci 50 kb away from each other should encounter each other every few minutes (every few seconds for loci 10 kb away from each other), which is much faster than the 30-60 minute time scale over which H2A phosphorylation occurs (see *Supplementary Information, Looping Model Derivation* for a description of this estimate).

The probability *P*_*loop*_ that the chromosome is in a looped state – i.e. the probability that the DSB-bound kinase is in contact with an H2A located *x* kilobases away – depends on the ratio of *x* to *l*, where *l* is the chromatin’s Kuhn length. For H2As very close to the break (*x* << *l*), the stiffness of the chromosome makes the formation of a loop highly improbable since the energy required to bend the chromatin into a small loop is prohibitive. For H2As far from the break site (*x* >> *l*), loops are unlikely to form since the kinase and target histones tend to occupy different regions of space. Loop formation is most probable in the regime of *x* ≈ *l*. Specifically, the *P*_*loop*_ (*x l*) is maximal at *x* = 1.7*l* (58). When the chromosome is in the looped state, the kinase phosphorylates the H2A at a rate *k*_*cat*_. We introduce the model parameter*φ*, which is the product of *k*_*cat*_ and factors that take into account how close to the H2A and in what orientations the kinase can be in order to phosphorylate the H2A. The looping model therefore has three parameters: *k*_*init*_, *l*, and *φ*.

#### 3D diffusion model

Following the arrival and activation of a kinase at the DSB, the kinase leaves the break site and then diffuses along some 3D trajectory, hitting H2As and phosphorylating them. An H2A near the break is likely to be contacted by the kinase because many of the possible trajectories go through this H2A. An H2A far from the break is less likely to be contacted because few trajectories go through this H2A – most trajectories go off to other parts of the nucleus far from this H2A. While this is clearly true for short trajectories that cross the nucleus only once, can this still apply to long trajectories? When the kinase continues diffusing, it presumably crosses the nucleus a large number of times over the course of the hour-long experiment, so in the later stages of its diffusion, the kinase is equally likely to hit any H2A in the nucleus. However, as there are so many H2As in the nucleus, it is reasonable to assume that the catalytic rate is only fast enough for the kinase to phosphorylate a small fraction of the H2As in the nucleus. Therefore, the later stages of a kinase’s diffusion might add only a negligible γ-H2AX signal. The predominant γ-H2AX signal comes from the early stage of diffusion, which results in the preferential phosphorylation of H2As near the break.

To predict the probability that the kinase will hit an H2A during the early stage of diffusion, we use the simple approximation *P*_*contact*_ = *a*/*R* (59), where *a* is the radius of the H2A and *R* is the 3D distance between the H2A and the kinase at the beginning of the kinase’s trajectory (*R* is taken to be the root mean squared distance from the H2A to the DSB, given by a worm-like chain treatment of the chromatin at thermodynamic equilibrium). We assume many copies of the kinase come on and off the DSB, where *k*_*leaveDSB*_ is the rate at which a kinase leaves the DSB: if kinases come off the DSB at a higher frequency, more kinases will be diffusing and contacting H2As, so H2A phosphorylation will occur at a faster rate. The 3D diffusion model has three parameters: *k*_*init*_, *l*, and *ω*, where *ω* is a product of *k*_*leaveDSB*_, *a* and other factors that describe how likely the kinase is to contact an H2A (see *Supplementary Information* for details).

#### 1D diffusion model

In the 1D diffusion model, the chromatin fiber is treated as a 1D filament of H2As. Starting from the break site, a kinase moves along the chromatin by sliding between adjacent H2A histones. At each step, the kinase phosphorylates H2A at a rate *k*_*cat*_. The kinase is allowed to move in the direction away from the break site or towards the break site with equal probability. The overall speed of the kinase’s motion is governed by the 1D diffusion coefficient *D*. For simplicity, we assume that the kinase does not permanently detach from the chromosome – i.e. we assume the time scale of falling off is greater than the duration of the experiment. Over a fixed time interval, the kinase is more likely to phosphorylate H2As that are closer to the DSB than those that are further because it takes a long time for a random walk to reach a distant location. The simulations include many copies of the kinase diffusing concurrently on the chromosome (this is further discussed in the *Supplementary Information*). The parameters of the 1D diffusion model are *k*_*init*_, *D*, and *k*_*cat*_.

#### Directed sliding model

As in the 1D diffusion model, the chromatin is treated as a 1D filament of H2A histones. Starting from the break site, the kinase slides along the chromatin at a rate *k*_*slide*_ until it falls off the chromosome at a rate *k*_*off*_. We assume the kinase phosphorylates all H2As along the way between the break site and wherever the kinase happens to fall off the chromosome, and the kinase cannot rebind to H2As that it has already phosphorylated (i.e. the kinase cannot backtrack); therefore, the kinase slides unidirectionally away from the break site. In this model, H2As close to the break are more likely to be phosphorylated because a kinase is unlikely to reach distant H2As before it falls off the chromatin. The parameters of the directed sliding model are *k*_*init*_, *k*_*slide*_, and *k*_*off*_.

## Supporting information

Dataset for Li, Bronk, Kondev and Haber

## Acknowledgments

We thank members of the Haber and Kondev groups and Dr. Douglas Theobald for helpful discussions. We also thank Dr. Maria Pia Longhese for providing us with plasmid PML105.45. Funding for JEH is provided by GM20056 and R35127029. Funding for JK is provided by the National Science Foundation grants DMR-1610737 and MRSEC-1420382, and by the Simons Foundation.

## Supplementary Information

### Supplementary Text

#### Model Derivations

All models are derived below and plotted in Figures S3-S14.

##### DSB Formation

All models begin with the formation of the DSB. We experimentally measure the cumulative probability distribution of the DSB formation time (Figure S1), and in all models we use this distribution by approximating it as a 12 minute lag time followed by the formation of the DSB at a rate of *k*_*DSB*_ = 0.08 minute.

#### Directed Sliding Model Derivation

##### Derivation Assuming a Kinase is at the Break Site at Time 0

We first derive a very simple model (in the next section we go on to a more detailed model, which is the one we compare to the experimental data). We model the chromosome as a 1D lattice of H2As: the H2As are at positions 1, 2, 3… and the break site is at position 0. The kinase is at position 0 at time 0 and then stochastically steps from one position to the next at a rate *k*_*slide*_. At time *t*, the probability that the kinase is at position *i* is 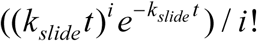. This is simply the Poisson distribution.

Next we include the process of the kinase falling off the chromosome at a rate *k*_*off*_, regardless of where on the chromosome the kinase is located. 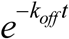 is the probability that the kinase is still on the chromosome at time *t. P*_*kinase*_ (*i,t*), the joint probability that the kinase is still on the chromosome AND is at the *i*-th H2A at time *t*, is therefore given by the product of the Poisson distribution and 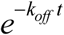:

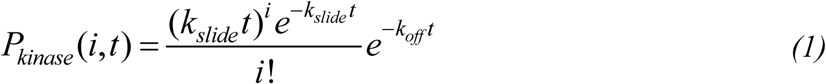

*P*_*phos*_ (*i,t*), the probability that the H2A at position *i* has been phosphorylated by time *t*, is given by

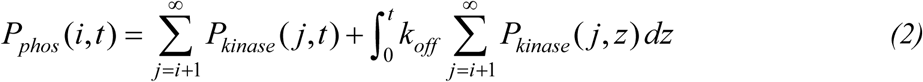

This formula follows from the assumption that the kinase phosphorylates every H2A that it comes to. Hence, if at time *t* the kinase is currently on an H2A at any position *j* > *i*, then we know that the kinase has already been to the *i*-th H2A and has already phosphorylated it. The first term in equation (2) is the probability that at time *t* the kinase is currently at any location beyond position *i*. (The summation goes to infinity for mathematical simplicity. Because *P*_*kinase*_ (*j,t*) is small at large *j*, there would be little change if we used a summation limit equal to the actual number of H2As in the chromosome). Similarly, if the kinase has fallen off the chromosome from any position *j > i* at any time before *t*, then we know that the *i*-th H2A has already been phosphorylated. The second term in equation (2) is the probability that the kinase has fallen off the chromosome from anywhere beyond position *i* at any time before time *t*.

Next we simplify the summation term in equation (2):

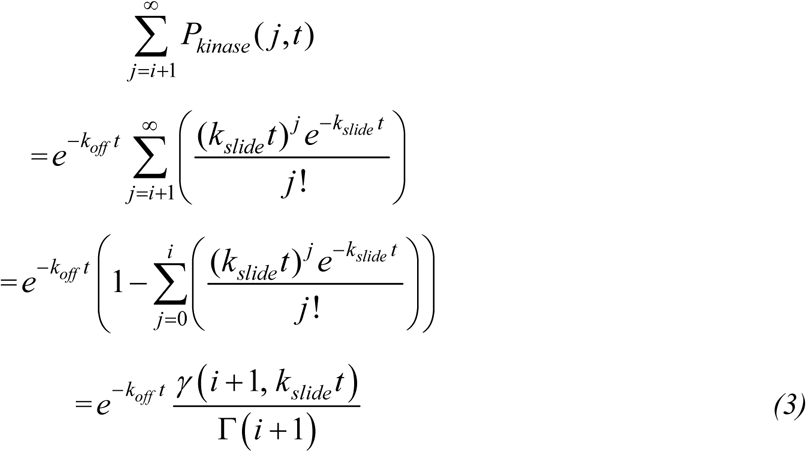

where Γ is the complete gamma function, and *γ* is the lower incomplete gamma function. Plugging expression (3) into equation (2), we obtain

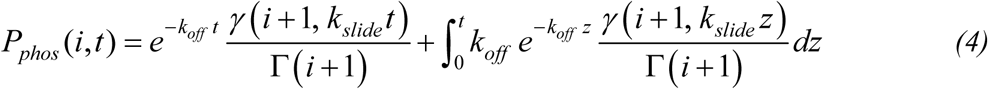

After evaluating the integral and simplifying, we obtain

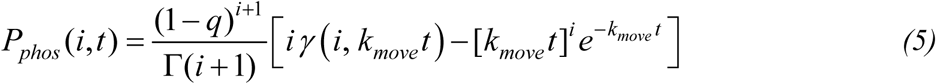

where 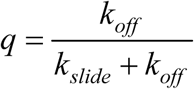 and *k*_*move*_ = *k*_*slide*_ + *k*_*off*_

#### Derivation that Incorporates the Variability in the Timing of DSB Formation and Arrival of the Kinase to the Break Site

Until this point, we have said that at *t* = 0 the kinase is at the break site and begins sliding along the chromosome. However, in different cells in the yeast population, the kinase starts sliding at different times (this will be detailed below). For an individual yeast cell, let *τ* be the time when the kinase begins the sliding process – i.e. the kinase does exactly what we described and derived so far, but beginning at time *τ* rather than time 0. Thus, we write*P*_*phos*_ (*i,t* | *τ*), which is the probability that the H2A at position *i* has been phosphorylated by time *t*, given that the kinase began the sliding process at time *τ*:

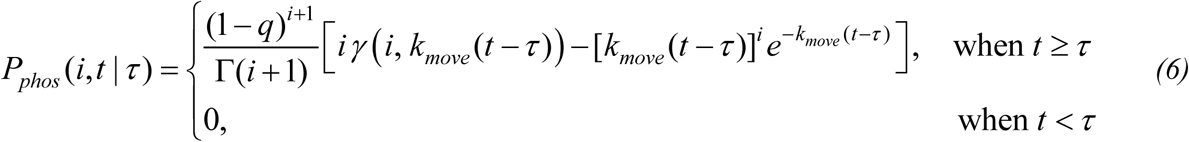

The sliding start time is stochastic and is described by the probability density function *PDF*(*τ*), which is derived in the following manner: The chromosome begins intact, which we call state I. The DSB forms at a rate *k*_*DSB*_: the chromosome is said to be in state II when there is a DSB but the kinase is not yet sliding. We make the assumption that once a yeast cell has a DSB, there is a single rate-limiting step described by the rate *k*_*init*_ that determines how soon the kinase initiates the sliding process (i.e. how soon the kinase arrives to the break site and then leaves the break site to slide along the chromosome). As soon as the kinase begins the sliding process, the chromosome is said to be in state III. Thus, *k*_*DSB*_ is the transition rate from state I to state II, and *k*_*init*_ is the transition rate from state II to III; this is captured by the coupled differential equations and initial conditions (7). (To simplify the derivation of the sliding model, in each yeast cell we say that only one copy of the kinase arrives at the break site; additional copies would have little effect on the predicted γ-H2AX profile because this derivation assumes that the kinase phosphorylates every H2A that it comes to). Let the functions *I(t), II(t)*, and *III(t)* represent the probabilities of being in states I, II, and III, respectively. We have

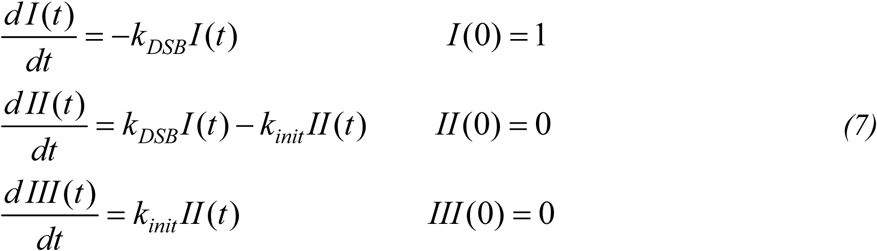

*PDF*(*τ*) is the probability per unit time that a cell will enter the state in which the kinase is sliding (i.e. state III). Therefore, 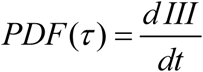. Solving equations (7) we obtain

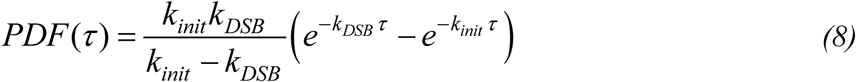

We calculate *P*_*phos*_ (*i,t*), the probability that the H2A at position *i* has been phosphorylated by time *t*, where the time when the kinase begins the sliding process is distributed according to equation (8). *P*_*phos*_ (*i, t*) is given by the integral of the product of equations (5) and (8):

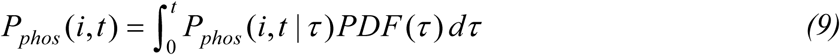

Evaluating this integral gives

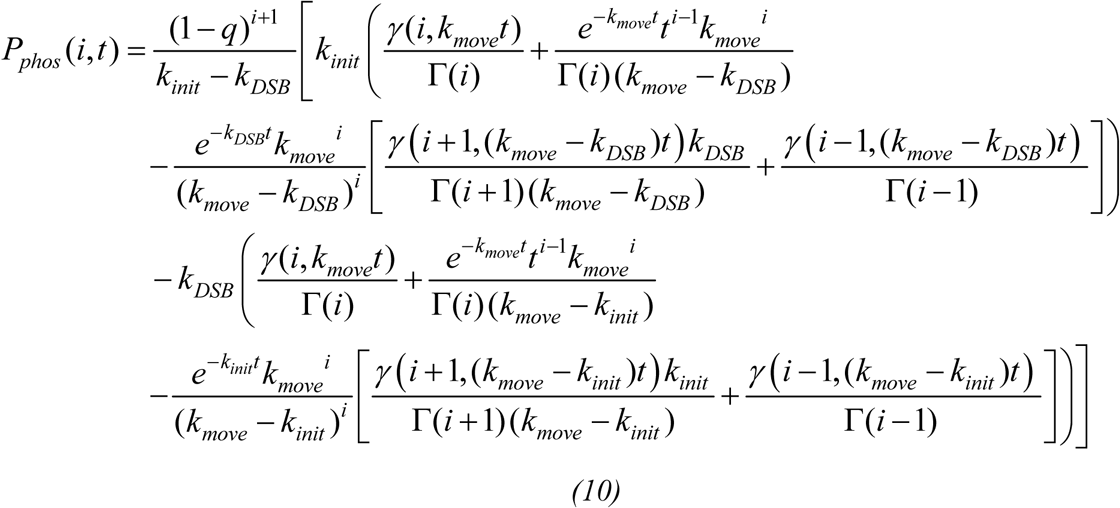

#### Looping Model Derivation

We model the chromosome as an unconfined worm-like chain at thermodynamic equilibrium. It is not necessary to include confinement by the nuclear envelope because we are only interested in the loci where phosphorylation occurs, which are 50 kb or less from the DSB. At thermodynamic equilibrium, there is a roughly 0.4 μm RMSD between two loci separated by 50 kb of chromatin (we estimated this based on the persistence length and compaction of the chromosome from Arbona et al. (1) and used the formula for *R* in the *Supplementary Information, 3D Diffusion Model Derivation*). The nucleus has a diameter of 2 μm, which is much larger than the 0.4 μm RMSD, so loci at 50 kb from the DSB are not confined by the nuclear envelope to stay in closer proximity to the DSB.

Using the equilibrium distribution of worm-like chain conformations is justified because we estimate that loci 50 kb away from each other should encounter each other every few minutes (every few seconds for loci 10 kb away from each other), which is much faster than the few tens-of-minutes time scale over which H2A phosphorylation occurs. We can estimate the contact frequency between two loci because we know the root mean squared distance *R* at equilibrium for the two loci. We also know the time *t* for a locus to diffuse a distance equal to *R* – this is obtained by inspecting the mean squared displacement (MSD) vs. time plot in Hajjoul et al. (2). Counterintuitively, *t* is equivalent to the time it takes for the two loci to come in contact. This is because chromatin undergoes subdiffusion, and any object moving subdiffusively thoroughly explores its space (3) – i.e. in the time *t*, the locus thoroughly explores a sphere of radius *R*. Therefore, loci within a distance *R* of each other will come in contact during the time *t*.

Now we introduce the formula for the worm-like chain looping probability. Let *l* be the Kuhn length of the worm-like chain and *x* be the contour length between two particular locations on the chain.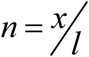 is therefore the number of Kuhn lengths between the two locations. *P*_*loop*_ is the fraction of chain conformations that have these two locations in physical contact. To be considered in physical contact, we say that the 3D distance between the two locations must be less than g, where g is a small fraction of a Kuhn length (g is unitless because it is a fraction). *P*_*loop*_ is given by

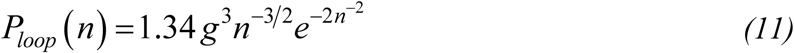

Note that equation (11) is an approximation – it does not predict *P*_*loop*_ exactly. Nevertheless, we compared equation (11) to worm-like chain simulations, and we found that the formula is accurate (i.e. has a maximum error of about 10%) for 1 ≤ *n* ≤ 20 and *g* ≤ 0.1. Equation (11) is based on ref. (4).

Now we incorporate the kinase into the model. Suppose the kinase is at the break site, and consider an H2A that is separated from the DSB by *n* Kuhn lengths of chromatin. *P*_*loop*_ (*n*) is therefore the probability that the chromosome is in a looped conformation in which the kinase and H2A are in physical contact. For H2A phosphorylation to occur, not only is this contact necessary but the kinase must also be in the necessary orientation. Let *ϵ* be the fraction of orientations that can result in phosphorylation, given that the H2A and kinase are in contact. Let *k*_*cat*_ be the rate at which the kinase phosphorylates the H2A, given that the kinase is in contact with the H2A and is in the necessary orientation. Therefore, the rate of phosphorylation of the H2A is

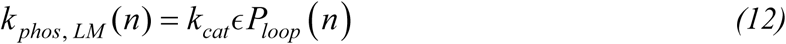

where the “LM” subscript stands for the looping model. Substitute equation (11) into equation (12), and replace *n* with 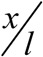, and we obtain

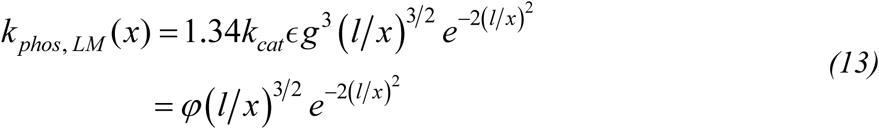

where *φ* = 1.34 *k*_*cat*_*ϵ g*^3^. Note that for *x* and *l* we use units of kilobase pairs because in our experiments we know the number of kilobase pairs between the DSB and our measured loci.

We construct the model to be consistent with ChIP measurements of Tel1 and Mec1 at the break site, which show their levels increase linearly with time (15, 29). These measurements could indicate two different dynamics. First, they could imply that there is an increasing number of kinases at the break site, and higher numbers of kinases could result in proportionally higher rates of H2A phosphorylation (this model was not considered in the main text). The phosphorylation rate is therefore the product of *k*_*phos, LM*_ (*x*) and the number of kinase copies on the DSB. We will refer to this as the “linearly increasing phosphorylation rate.”

Alternatively, even with multiple copies of kinases bound near the break site, it is possible that only one or two copies perform phosphorylation. (Hypothetical scenarios that would cause this include these possible examples: there is a row of kinases near the break site, but only the kinase precisely at the DSB is activated, or steric hinderance causes only the kinase copies at the ends of the row to have their active sites accessible to H2As). Thus, in each cell, once a kinase has arrived to the DSB, there is simply a constant rate of H2A phosphorylation given by *k*_*phos, LM*_ (*x*) (this model was considered in the main text). We will refer to this as the “constant phosphorylation rate.” The constant and linearly increasing phosphorylation rates are the two extremes in possible dynamics; intermediate dynamics could occur – i.e. phosphorylation rates that increase in a sub-linear manner. We only examine the two extremes because they yield the greatest difference in predicted γ-H2AX profiles. We will begin with a derivation that involves the constant phosphorylation rate.

#### Looping Model with a Constant Phosphorylation Rate

This is the model discussed in the main text.

When a kinase is present on the break site, it phosphorylates H2As at a rate *k*_*phos, LM*_ (*x*), so we have

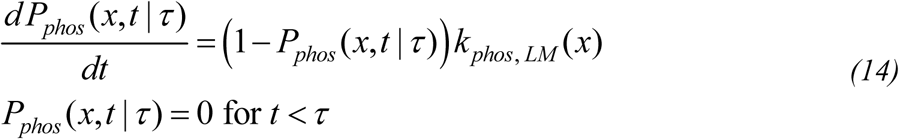

where *P*_*phos*_ (*x,t* |*τ*) is the probability that the H2A at *x* kilobase pairs from the break site has been phosphorylated by time *t*, given that the kinase arrived to the break site at time *τ*. (The bottom equation of equations (14) states that no phosphorylation takes place until the kinase arrives to the break site). Solving equations (14) gives

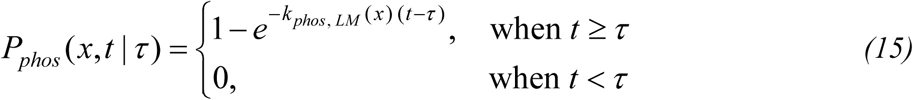

Recall that in the “constant phosphorylation rate” version of the looping model, we only consider a single copy of the kinase at the break site because we assume that additional copies of the kinase would not change the phosphorylation rate. In different cells in the yeast population, the single copy of the kinase arrives to the break site at different times. To take this into account, we perform the same derivation that is in the directed sliding model’s section entitled “Derivation that Incorporates the Variability in the Timing of DSB Formation and Arrival of the Kinase to the Break Site:” *P*_*phos*_ (*x,t*), the probability that the H2A at *x* kilobase pairs from the break site has been phosphorylated by time *t*, is given by

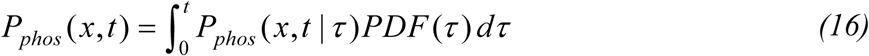

where *P*_*phos*_ (*x,t* |*τ*) is given by equation (15) and *PDF*(*τ*) is given by equation (8). Note that when using equation (8) for the looping model, *k*_*init*_ indicates the rate at which the kinase arrives to the break site (note that this differs from the definition of *k*_*init*_ used in the sliding model because in the sliding model *k*_*init*_ involves kinase arrival to the break site AND the process of leaving the break site to begin sliding). Upon carrying out the integration in equation (16), we obtain

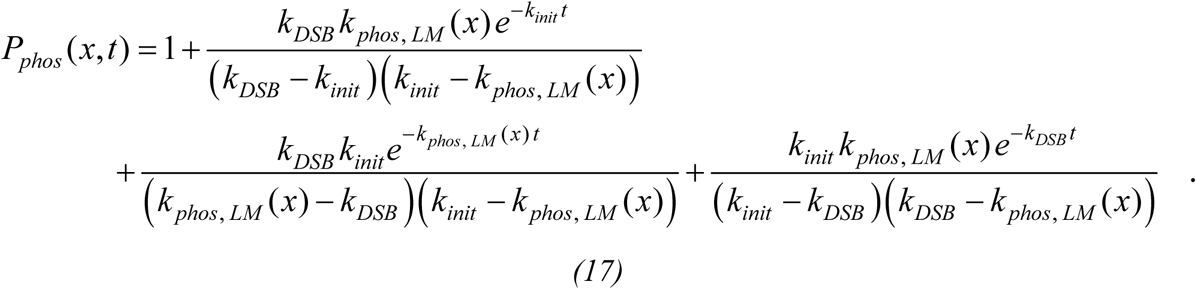

#### Looping Model with a Linearly Increasing Phosphorylation Rate

This model is not included in the main text because the Bayes factor calculations demonstrate that it is worse than the model with the constant phosphorylation rate. In this version of the looping model, the phosphorylation rate is proportional to the number of copies of the kinase that are on the break site. Let *M* (*t* | *δ*) be the average number of copies of the kinase on the break site at time t, given that the DSB formed at time *δ*. Therefore, we have

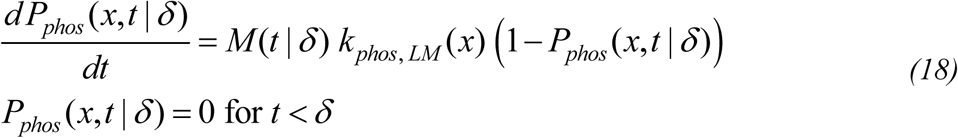

where *P*_*phos*_ (*x,t* | *δ*) is the probability that the H2A at *x* kilobase pairs from the break site has been phosphorylated by time *t*, given that the DSB formed at time *δ*. (For simplicity, we obtain an approximate solution by using the mean number of kinase copies on the break site *M* (*t* | *δ*), ignoring the variance in the numbers of kinase copies.)

The average number of kinase copies at the break site is linear with time (i.e. kinases arrive with a constant rate *k*_*init*_), so *M* (*t* | *δ*) = (*t* – *δ*)*k*_*init*_. Solving equations (18) using this expression for *M*(*t* | *δ*) gives

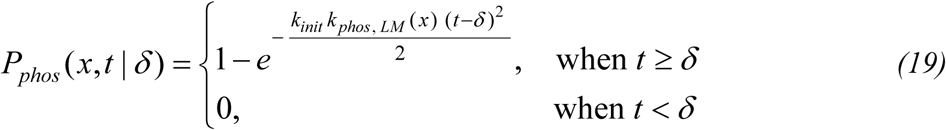

Next we take into account that *δ* varies across yeast cells in the population. Because the DSB forms at a constant rate *k*_*DSB*_, there is an exponential distribution of waiting times for DSB formation, given by the following probability density function for *δ*:

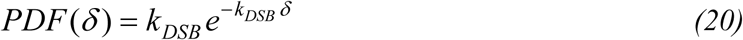

*P*_*phos*_ (*x,t*), the probability that the H2A at *x* kilobase pairs from the break site has been phosphorylated by time *t*, is given by

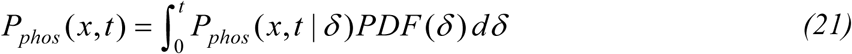

We plug equations (19) and (20) into equation (21) and evaluate the integral numerically.

Note that in equation (19), *k*_*init*_ and *k*_*phos, LM*_ (*x*) never appear separately: they are always multiplied together. Using equation (13), we have 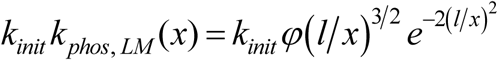, so we define the parameter *ψ* = *k*_*init*_*φ* (i.e. *k*_*init*_ and *φ* are not separate parameters in the “Linearly Increasing Phosphorylation Rate” version of the looping model). Therefore, this version of the looping model has only two parameters: *ψ*, and *l*.

#### 1D Diffusion Model Simulation

Here we describe additional details of the Gillespie simulations for 1D diffusion that were not discussed in the *Computational Methods* section of the main text. As previously mentioned, we construct all of our models to be consistent with ChIP measurements of Mec1 and Tel1 at the break site, which show Mec1 and Tel1 levels that increase linearly with time. In the 1D diffusion model, linearly increasing kinase levels could be achieved through one of two simple mechanisms, one in which there is a constant rate at which kinases initiate 1D diffusion, and a mechanism in which the rate of diffusion initiation increases linearly with time. The former mechanism is discussed in the main text, while the latter mechanism is not since our Bayes factor calculation demonstrates that the former is more likely to be correct. We will now describe how these mechanisms give rise to linearly increasing kinase levels at the break site and will discuss how these mechanisms are implemented in our simulations.

We begin by describing the mechanism with a constant rate of diffusion initiation. Let *k*_*arriveDSB*_ be the rate at which a kinase arrives to the break site and *k*_*leaveDSB*_ be the rate at which a kinase leaves the break site to start diffusing along the chromosome. In the mechanism with a constant rate of diffusion initiation, only one kinase copy can leave the break site at a time. (This could happen if the copies of the kinase line up in a row close to the break site, such that the one copy furthest from the DSB is physically blocking the other copies from being able to slide away from the break site). In this scenario, if *k*_*arriveDSB*_ > *k*_*leaveDSB*_, then copies of the kinase will build up at the break site linearly with time: we will have *M* (*t*) = (*k*_*arriveDSB*_ – *k*_*leaveDSB*_)(*t* – *δ*), where *δ* is the time when the DSB forms, and *M* (*t*) is the average number of kinases on the break site (*t* – *δ*) minutes after the formation of the DSB.

We perform this simulation by having a single rate-limiting step described by the rate *k*_*init*_, where *k*_*init*_ is the rate at which a kinase copy arrives to the break site and then departs the break site to diffuse along the chromosome. (Note that it is only necessary to use this single rate rather than the two rates *k*_*arriveDSB*_ and *k*_*leaveDSB*_: using the single rate becomes equivalent to using two rates if *k*_*arriveDSB*_ is fast. If *k*_*arriveDSB*_ is slow, then the situation becomes equivalent to the mechanism in which the rate of diffusion initiation increases linearly with time, which is discussed in the next paragraph.) Throughout the entire duration of the simulation, kinase copies continue to start at rate *k*_*init*_. Therefore, in our simulations there can be many kinase copies diffusing concurrently on the chromosome.

In the mechanism in which the rate of diffusion initiation increases linearly with time, the kinases do not block each other, but rather all kinases on the break site are capable of sliding away from the break site. In this case, the number of kinases that leave the break site per unit time is *M* (*t*) *k*_*leaveDSB*_. If *k*_*arriveDSB*_ >> *M* (*t*) *k*_*leaveDSB*_, then copies of the kinase will build up at the break site linearly with time: we will have *M* (*t*) = *k*_*arriveDSB*_ (*t* – *δ*). The number of kinases that leave the break site per unit time will therefore be *k*_*leaveDSB*_ *k*_*arriveDSB*_ (*t* – *δ*). In other words, unlike in the first mechanism where there is a constant rate of kinase copies starting to diffuse along the chromosome, in this mechanism the rate of kinase copies starting to diffuse grows linearly with time. We implement this mechanism’s simulation by having each kinase copy arrive to the break site at a rate *k*_*arriveDSB*_ and leave the break site at a rate *k*_*leaveDSB*_. Note, however, that these rates are not separate parameters because the dynamics are governed by the product *k*_*leaveDSB*_ *k*_*arriveDSB*_, so when we vary parameter sets, we only vary *k*_*arriveDSB*_, and we simply set *k*_*leaveDSB*_ = 0.0005 /minute (we chose this value since it is slow enough for there to be linear growth in the number of kinases at the break site over several hours). When we report parameter values, we report *z*, where *z* = *k*_*leaveDSB*_ *k*_*arriveDSB*_. Kinase copies begin diffusing throughout the entire duration of the simulation, so there can be many kinase copies diffusing concurrently on the chromosome. For simplicity, in all 1D diffusion simulations we assume the kinases do not interact with each other (i.e. no traffic jams); to avoid the regime where traffic jams would become likely, we limit how large *k*_*init*_ and *z* can be in order to limit the total number of kinases along the chromosome.

#### 3D Diffusion Model Derivation

We treat the kinase as a randomly diffusing particle and each H2A histone as a spherical target of radius *a*. Given infinite time, a particle diffusing without constraints can either hit the target or diffuse infinitely far away without ever hitting the target – the probability that the particle hits the target is *P*_*contact*_ = *a /R* (7), where *R* is the 3D distance between the target and particle at the beginning of the particle’s trajectory. The infinite time assumption approximates our situation because the hour time scale of phosphorylation is much longer than the seconds time scale presumably required for proteins to diffuse across the nucleus (see ref. (8) for diffusion in the nucleus of human cells). The assumption of no constraints can be applied to our system despite the presence of the nuclear envelope. Confinement by the nuclear membrane keeps the kinases in the nucleus, so the kinases presumably continue phosphorylating H2As, resulting in a background γ-H2AX signal that is approximately the same for all H2As. It is a reasonable assumption that the kinase’s catalytic rate is slow enough that over the time scale of the experiment, the kinases would only have time to phosphorylate a small fraction of the H2As throughout the nucleus. The background signal would therefore be minimal and can be ignored, allowing us to simply have *P*_*contact*_ = *a*/*R*. (Alternatively, it is also possible that the background signal is not negligible, but having a significant background signal would make the 3D diffusion model not match the experimental γ-H2AX profiles, so we do not consider this scenario).

The kinase starts its trajectory at the break site, and the target is an H2A located *x* kb from the DSB. The distance *R* is taken to be the root mean squared distance from the H2A to the DSB, given by a worm-like chain treatment of the chromatin at thermodynamic equilibrium: 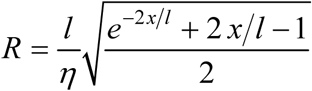, where *l* is the chromatin’s Kuhn length, and *η* is the chromatin compaction, the number of kilobases of DNA per nanometer of chromatin fiber. Therefore, 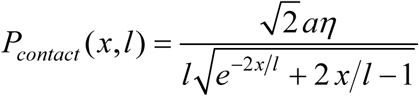. According to *P*_*contact*_, H2As further from the DSB are less likely to be contacted by the kinase.

We assume many copies of the kinase come on and off the DSB, where the rate of leaving the DSB is *k*_*leaveDSB*_. Therefore, an H2A located *x* kb from the DSB is hit by kinases at a rate *P*_*contact*_ (*x,l*)*k*_*leaveDSB*_. If *υ* is the fraction of contacts that results in phosphorylation, the rate at which the H2A is phosphorylated is *k*_*phos*, 3*DDM*_ (*x*) = *υP*_*contact*_ (*x,l*)*k*_*leaveDSB*_ (the subscript “3DDM” stands for the 3D diffusion model). Plugging in the expression for *P*_*contact*_, we have

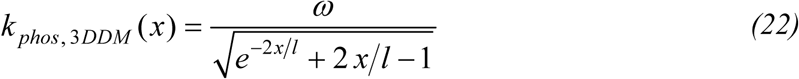

where 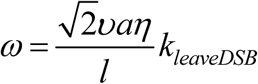.

There are two versions of the 3D diffusion model: one with a constant rate of diffusion initiation (which is the model discussed in the main text), and a version with a linearly increasing rate of diffusion initiation. The rationale for trying these two model variants is the same as that described in the *1D Diffusion Model Simulation* section above. The Bayes factor analysis determined that the 3D diffusion model with the constant rate is more likely to be correct than the 3D diffusion model with the linearly increasing rate.

The 3D diffusion model with a constant rate of diffusion initiation involves kinases leaving the break site at a constant rate and therefore phosphorylating H2As at a constant rate. This model is derived in exactly the same way as the looping model with a constant phosphorylation rate (equation (17)), except the phosphorylation rate *k*_*phos, LM*_ (*x*) is replaced by *k*_*phos*, 3*DDM*_ (*x*), so we obtain

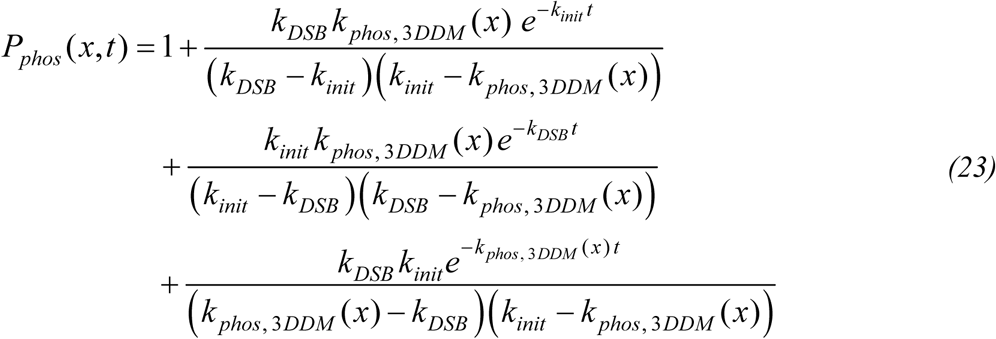

The 3D diffusion model with a linearly increasing rate of diffusion initiation involves kinases leaving the break site at an increasing rate. This model is derived in exactly the same way as the looping model with a linearly increasing phosphorylation rate (equations (19)-(21)), except the phosphorylation rate *k*_*phos, LM*_ (*x*) is replaced by *k*_*phos*, 3*DDM*_ (*x*), so we have the following equations:

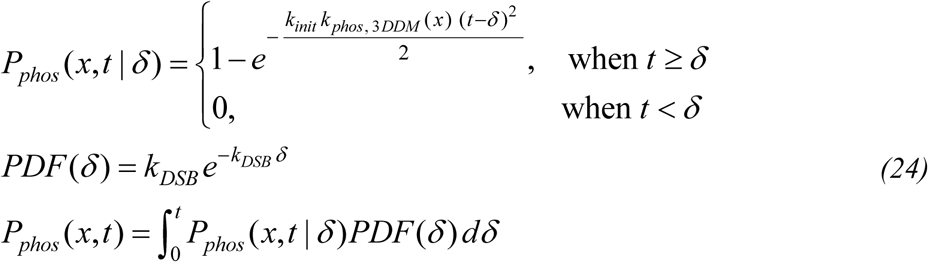

where the integral is evaluated numerically. *k*_*init*_ and *k* _*phos*, 3 *DDM*_ always appear multiplied together. From equation (22), we see that 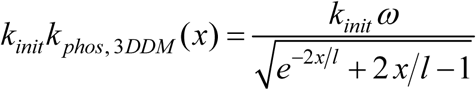, so we define the parameter *ζ= k*_*init*_ *ω*. Therefore, this version of the 3D diffusion model has only two parameters: *ζ*, and *l*.

#### Modeling Phosphorylation Spreading Around the Recombination Enhancer

When predicting γ-H2AX levels near the Recombination Enhancer (RE), we take into account the protein-mediated binding of RE to the break site at the *MAT* locus (23, 24). With RE bound to the DSB, RE-adjacent loci (that are 5 or 10 kb from RE) are now in close spatial proximity to the DSB, so the 3D models predict that the kinases can reach the RE-adjacent sites. 1D models predict that the kinases cannot reach RE-adjacent sites because RE is located 171 kb away from *MAT*. For 3D models, we include two additional parameters in order to take into account the binding of RE to *MAT*. Initially there is a period when RE is not bound to the DSB at all, but then proteins at *MAT* undergo phosphorylation at their threonine residues, which mediates the RE-*MAT* interaction. Therefore, we include in the model that at rate *k*_*RE*_ the cell transitions to the state in which RE and the DSB are bound some fraction of the time (denoted by *F*_*RE*_), at which point the kinase can phosphorylate RE-adjacent sites. The rate of phosphorylation for an H2A that is *x* kb from RE is assumed to equal the phosphorylation rate for an H2A that is *x* kb from the DSB multiplied by *F*_*RE*_. In other words, for the RE-adjacent sites, *P*_*contact*_ (*x,l*) in the 3D diffusion model and *P*_*loop*_(*x,l*) in the looping model are multiplied by *F*_*RE*_ because the kinase only comes in contact with the RE-adjacent sites during the fraction of the time when RE and *MAT* are bound. The optimal values of the parameters are *F*_*RE*_ = 0.19 and *k*_*RE*_ = 0.025/minute, which are consistent with the kinetics of mating type switching as measured in Avşaroğlu et al. (10). Because *k*_*RE*_ is subtantially slower than *k*_*init*_, *k*_*RE*_ is rate-limiting, so we simply treat *k*_*init*_ as equal to 0.025/minute in equations (17, 23).

#### Some Models Require Kinase Activation at the DSB

The 3D diffusion, 1D diffusion and directed sliding models require that the kinase becomes activated upon arrival to the break site. Without this activation, these models would not predict preferential phosphorylation of H2As close to the break site; the kinase becomes activated at the DSB and is more likely to encounter closer H2As than farther H2As. To further argue this point, consider a model in which the kinase is activated without having to encounter the DSB. At the moment when the DSB is formed, the kinase is at some arbitrary location in the nucleus, so it is equally likely to diffuse to any genetic locus and then start phosphorylating H2As from that locus (this applies for directed sliding, 1D diffusion and 3D diffusion). Simply assuming strong binding between the kinase and the break site would not result in increased phosphorylation near the DSB. Strong binding to the DSB would merely hold the kinase in place for a while – i.e. kinases that happen to come in contact with the DSB would end up wasting their time sitting on the DSB, so they would have less time to phosphorylate H2As. As previously noted, a precedent for kinase activation was found by an *in vivo* study of ATM, the mammalian homolog of Tel1: upon binding to the MRN complex at the break site, ATM undergoes autophosphorylation and thus becomes active (11).

Note that the 1D diffusion and directed sliding models are mathematically equivalent to models in which many copies of the kinase bind along the chromatin, each binding to the next adjacent H2A. In this case, kinase activation could take place but is not required in the model. This is because the strong binding of the kinase to the break site could serve to nucleate the formation of the row of kinases. The directed sliding model is also equivalent to a model in which the kinase remains bound to the break site while the nearby chromatin slides past it – e.g. this would happen if the cohesin complex binds to the DSB and extrudes chromatin through the cohesin ring, causing the DSB-bound kinase to slide past the nearby chromatin; such a model does not require kinase activation at the break site (chromatin extrusion by cohesin has been found to occur in mammals (12), though studies have not investigated whether it leads to the spreading of histone modifications).

Also note that the looping model does not require activation at the DSB: H2As are preferentially phosphorylated in the vicinity of the DSB because the kinase binds the DSB and the looping of the chromosome brings the kinase into contact with nearby H2As.

#### A quantitative model of ChIP

In order to quantitatively compare our phosphorylation propagation models to ChIP data, we must have a quantitative understanding of ChIP. It is not sufficient to merely have the qualitative understanding that a higher ChIP signal indicates the presence of more γ-H2AXs. Our phosphorylation propagation models predict the probability *P*_*phos*_(*x,t*) that an H2A has been phosphorylated, where *x* is the number of kilobase pairs separating the H2A from the break site, and *t* is the elapsed time since the addition of galactose (galactose induces production of HO-endonuclease, the enzyme that forms the DSB). Measuring the ChIP signal does not directly give us the probability of phosphorylation. Rather, the ChIP signal is some function of this probability: the function *S* (*P*_*phos*_) will represent the ChIP signal. Below, we make a model of the ChIP process in order to derive the functional form of *S*. Having obtained the functional form of *S*, we can use it in conjunction with the *P*_*phos*_(*x,t*) predicted by each phosphorylation propagation mechanism: We plug in the predicted *P*_*phos*_(*x,t*) as the argument of *S* to yield the predicted γ-H2AX ChIP profile *S*(*P*_*phos*_ (*x,t*)) for each mechanism of phosphorylation propagation. We can then compare each mechanism’s predicted γ-H2AX ChIP profile to the experimentally-measured ChIP profile.

The experimental ChIP signal that we report is 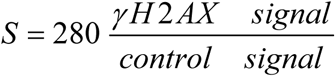. *γ-H2AX signal* is the qPCR signal for the DNA that has γ-H2AX bound to it; *control signal* is the qPCR signal from all DNA at the same locus (i.e. all DNA regardless of whether it has γ-H2AX on it). The 280 is present because the sample for the control was diluted by a factor of 280 relative to the sample for the *γ-H2AX signal*. We quantitatively model *S* by taking into account that only some of the DNA with γ-H2AX on it is pulled down by antibodies during ChIP: let *P*_*pull*–*down*_ (*P*_*phos*_) be the probability of pulling down DNA at the measured locus, which depends on the *P*_*phos*_ at that locus (discussed further below). We also take into account that aside from the pull-down, other steps (e.g. washing steps) in ChIP may reduce the recovery of DNA. Let *C* be 280 multiplied the probability that the DNA will be recovered in all the other steps, given that it was pulled down by the antibodies. Therefore, our model for the ChIP signal is *S*(*P*_*phos*_) =*CP*_*pull*–*down*_ (*P*_*phos*_).

Let us now quantitatively model *P*_*pull*–*down*_. In the ChIP protocol, sonication results in DNA fragments roughly 500 bp in length, containing multiple H2As. The DNA fragment comes to the surface or interior of a porous agarose bead, where the DNA fragment’s γ-H2AXs can bind to antibodies that are attached to the bead. We make two assumptions: (1) The binding of a single antibody to a single γ-H2AX results in the DNA fragment being pulled down. (Even if there are multiple γ-H2AXs on a DNA fragment, only one γ-H2AX needs to be bound by an antibody for the DNA fragment to be pulled down). Let the parameter *f* be the probability that a particular γ-H2AX is bound by an antibody. (2) The second assumption is that the antibodies bind independently of each other – i.e. an antibody binding to one γ-H2AX has no effect on whether an antibody binds to another γ-H2AX on the same DNA fragment.

These two assumptions are captured mathematically in the following manner: Let *N*_*H*2*A*_ be the average number of phosphorylatable H2A’s on a DNA fragment – e.g. *N*_*H*2*A*_ =6 in cells with all wild-type H2As, and we use *N*_*H*2*A*_ =3 in cells with only 1/2 wild-type H2As. *N*_*H*2*A*_*P*_*phos*_ is the average number of γ-H2AXs on a DNA fragment. The probability that the DNA fragment is pulled down is 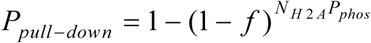 because this is the probability that at least one γ-H2AX is successfully bound by an antibody (i.e. it is 1 minus the probability that none of the γ-H2AXs are bound by antibodies). This expression is not valid for *N*_*H*2*A*_*P*_*phos*_ < 1 (i.e. if the average number of γ-H2AXs on a DNA fragment is less than 1). If *N*_*H*2*A*_*P*_*phos*_ < 1, then we use the expression *P*_*pull*–*down*_ = *N*_*H*2*A*_*P*_*phos*_ *f* because *N*_*H*2*A*_*P*_*phos*_ is the probability of one γ-H2AX being present on the DNA fragment, and *f* is the probability that the DNA fragment is pulled down given that there is one γ-H2AX on the fragment. Therefore, our quantitative model of the ChIP signal is the following:

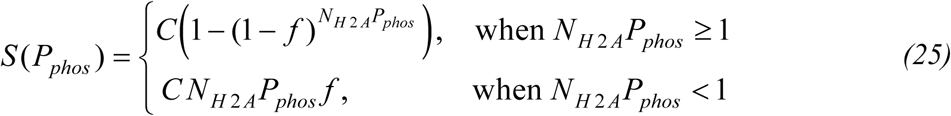

where *N*_*H2A*_, *f* and *C* are parameters of the quantitative ChIP model, and *P*_*phos*_ includes both the predicted DSB-dependent phosphorylation and the basal phosphorylation that occurs prior to DSB formation (see the next section on basal γ-H2AX levels). equations (25) are applicable to all models besides the directed sliding model, which is discussed below in *A quantitative model of ChIP for the directed sliding model*.

As an example of the predictions from this model of ChIP, suppose it were the case that all the phosphorylatable H2As at a locus were phosphorylated – i.e. *P*_*phos*_ (*x,t*) = 1. We use *f* = 0.16 and *N*_*H2A*_ = 6 for the wild type and *N*_*H2A*_ = 3 for the mutant strain, which are the best parameter values extracted from Bayesian parameter estimation (as shown in Table 2). Then although the mutant strain has only 1/2 as many γ-H2AXs as the wild type, the mutant strain will have a ChIP signal that is 0.63 times the signal of the wild-type strain.

In the looping and 3D diffusion models, phosphorylation of one H2A is independent of phosphorylation of another H2A, even in the case of H2As on the same DNA fragment. For the looping model, this is because phosphorylation occurs according to the thermodynamic equilibrium looping probabilities, so the phosphorylation of one H2A has no influence on whether a nearby H2A is phosphorylated. For the 3D diffusion model, a single copy of a kinase is unlikely to contact more than one H2A on a given DNA fragment since the probability of the kinase contacting an H2A is at most 0.1 (this is estimated by plugging in reasonable values for the coefficients in the formula for *P*_*contact*_ (*x,l*) in the *Supplementary Information, 3D Diffusion Model Derivation*); therefore, the phosphorylation of H2As is effectively uncorrelated. The independence of H2A phosphorylation means that the number of γ-H2AXs on a DNA fragment is binomially distributed with a peak at *N*_*H*2*A*_*P*_*phos*_. In the looping and 3D diffusion models, for simplicity we ignore the spread of the binomial distribution and just use *N*_*H* 2 *A*_*P*_*phos*_ as the number of γ-H2Axs on a DNA fragment. This simplifying assumption is what we included in equations (25).

In the 1D diffusion model, however, phosphorylation of one H2A is not independent of another H2A on the same DNA fragment, and so it is possible that there would be a wide range in the number of γ-H2AXs on a DNA fragment, and so using just the mean number may not be accurate. Therefore, in every run of the 1D diffusion simulation, we record the number of γ-H2AXs on each DNA fragment. We apply the ChIP model to every DNA fragment and then take the average over the fragments from all runs in order to get the net ChIP signal.

#### Basal γ-H2AX levels are incorporated into the predicted ChIP signals

Experimentally we measure a basal level of γ-H2AX on the chromosome that exists at time 0 (which is before the DSB forms) – this is the ChIP signal measured at time 0 in Figures 1, 2, 3, 5, 6 and 7. We use the mean values of the basal γ-H2AX profile to serve as the initial ChIP signal in every model of γ-H2AX spreading (rather than starting from a ChIP signal of 0). Because basal phosphorylation and DSB-dependent phosphorylation are presumably independent of each other, the probability of phosphorylation as a result of either basal or DSB-dependent phosphorylation is

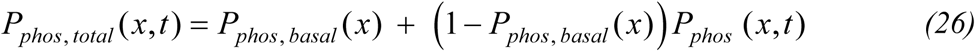

where *P*_*phos*_ (*x,t*) is the DSB-dependent phosphorylation derived for the model of γ-H2AX spreading. To obtain *P*_*phos, basal*_, we note that the basal ChIP signals are small, so *P*_*phos, basal*_ must be small. Therefore, we can use the bottom equation (25), and rearrange to get 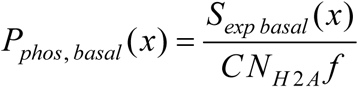, where *S*_*exp basal*_ (*x*) is the experimentally measured basal ChIP signal. To obtain the predicted ChIP signals, *P*_*phos, total*_ (*x,t*) is what we actually use in equations (25) in place of *P*_*phos*_

#### A quantitative model of ChIP for the directed sliding model

For the directed sliding model, we modify equations (25). In the directed sliding model we assume that whenever a kinase slides to a locus, it phosphorylates all H2As there. However, the kinase does not make it all the way to every locus. In the directed sliding model, *P*_*phos*_ (*x,t*) is equivalent to the probability that the kinase arrives to location *x*. If the kinase does not arrive to the location *x*, then the ChIP signal is the basal ChIP signal *S*_*exp basal*_ (*x*), but if the kinase does arrive to the location, then the ChIP signal is 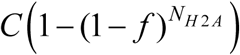. Therefore, the ChIP signal for the directed sliding model is given by

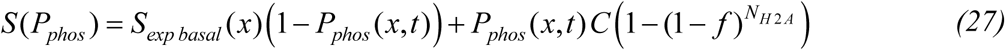

#### Bayes factor calculation

The Bayes factor is given by the ratio of marginal likelihoods for two models, Models A and B:

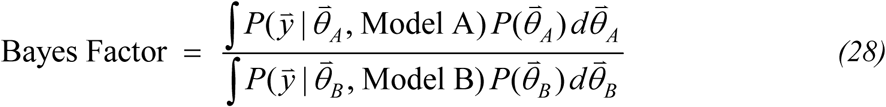

where *P*(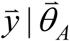, Model A) is the likelihood of the experimental data 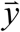 given Model A and the model parameters 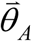, and 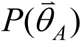 is the prior probability of 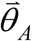 (likewise for Model B). These are discussed below. In the tables displaying the Bayes factors (Table 1 and S1), Model B is always the model with the highest marginal likelihood compared to the other models in the table.

We use previous studies to estimate the range of possible parameter values – the integral in equation (28) is over this range. The previous studies measured the chromatin Kuhn length (95% CI is 8.4-15 kb) and linear density of chromatin DNA (95% CI 56-68 bp/nm) (1), and density of the nucleosomes along the chromatin (about 170 bp/nucleosome) (12) and provided data on the kinetics of *in vivo* H2A phosphorylation that allow us to constrain kinetic parameter ranges (12). Ref. (10) was used to constrain RE parameters. In some cases, there is some arbitrariness in the choice of the range, but choosing a different range (that is still reasonable given the previous studies) only changes the Bayes factors by an order of magnitude. The ChIP parameter ranges are as follows: *f* is allowed to vary between 0 and 1, N is allowed to be 2, 4 or 6 H2As, and *C* is allowed to vary between 8.3 and 280, which is based on our measurements of DNA recovery from ChIP compared to samples in which PCR was performed without ChIP.

The functional form of the prior probability distribution *P*(*θ*) for each parameter *θ* is as follows: For the parameters whose values we know within an order of magnitude (i.e. the Kuhn length *l* and ChIP parameter *N*_*H*2*A*_), we use the uniform prior *P*(*θ*) = constant. For parameters that are less certain, we use Jeffreys priors. However, it is difficult to obtain exact Jeffreys priors given the complexity of our models’ functional forms, so we use Jeffreys priors from similar but simpler scenarios. For parameters that represent the probability of a binary event (i.e. the ChIP parameter *f*, the directed sliding model’s parameter *q*, and the RE-MAT binding fraction *F*_*RE*_), 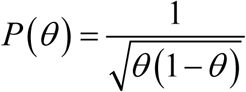 (16). All other parameters are assigned prior distributions of 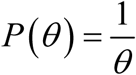, which is the Jeffreys prior for the rate parameter of an exponential distribution. All prior distributions are normalized and constrained to be within the estimated range of possible values (i.e. *P*(*θ*) = 0 if *θ* is outside the range of possible values). For simplicity, we assume that 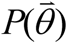 for the parameter set 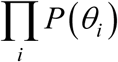 is the product of the priors 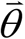 for the individual parameters *θ*_*i*_.

Because our priors are not precisely Jeffreys priors, let us estimate the extent to which this lack of precision could affect the Bayes factor. To do so, one can calculate the Bayes factor using a different prior. For example, consider using a uniform prior for all model parameters for both the directed sliding model and the 1D diffusion model. For most of the parameters, a uniform prior is quite different from the Jeffreys prior; the uniform prior will serve as an extreme example, so the Bayes factor will presumably be changed by more than it would be if we were able to use the exact Jeffreys prior. As shown in Table 1*B* of the main text, the Bayes factor is 10^−6^ when Mec1 is undergoing 3D diffusion and Tel1 is undergoing 1D diffusion (this Bayes factor is in comparison to Mec1 3D diffusion with Tel1 directed sliding). Using all uniform priors, this Bayes factor becomes 10^−7^. Therefore, using a prior with a very different functional form can change the Bayes factor by an order of magnitude but does not change our conclusions.

Finally, we discuss the likelihood function. Consider a particular theory, e.g. Model A. For a particular parameter set 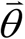 and for the *i*th measurement condition (each measurement condition is composed of the yeast strain, the time point and the location on the chromosome), Model A predicts a single γ-H2AX ChIP signal *S*_*i*_. However, because there is measurement error in the experiments, our theory should predict that the experimental data will have some spread about the mean predicted ChIP signal. Therefore, we incorporate an error model into the theory, which we assume to be Gaussian error – i.e. model A predicts that the mean ChIP signal is *S*_*i*_ and predicts a Gaussian spread about the mean with standard deviation ***σ***_*i*_. Hence the likelihood function is

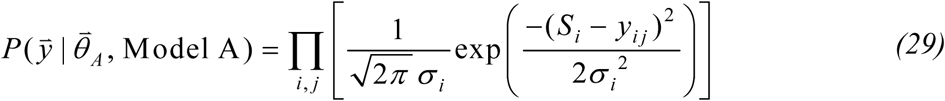

The experimentally measured ChIP signal *y*_*ij*_ has the subscript *i* for the measurement condition and the subscript *j* because there are multiple data collected for the same measurement condition.

Analysis of all of our ChIP data revealed that for each measurement condition, the standard deviation of the ChIP data is approximately equal to the mean ChIP signal multiplied by 0.35. Therefore, our error model can use ***σ***_*i*_ = 0.35*S*_*i*_. Including this into equation (29) yields

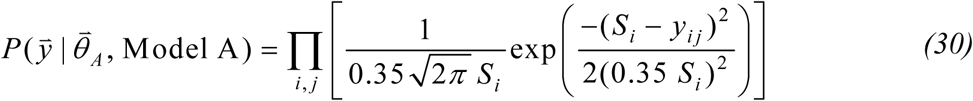

Finally, the integral in equation (28) is carried out by evaluating the integrand at points on a grid in parameter space and then using a Riemann sum.

As previously discussed, most theories have two variants: (1) a model in which there is an increasing phosphorylation rate or increasing rate of diffusion initiation, and (2) a model in which there is a constant phosphorylation rate or constant rate of diffusion initiation. For each theory, we use the best variant when reporting the Bayes factors. (We found that the constant phosphorylation rate/constant diffusion initiation rate models always fit the data better than the models with the increasing rates).

When calculating the Bayes factor for Tel1 and Mec1 models simultaneously, we exclude the Tel1 3D models because they were proven to be very unlikely from the Bayes factor analysis of Tel1 individually. This exclusion is justified because the analysis in Table 1*A* indicates that the 1D models are extremely favored for Tel1, and by inspecting the best ChIP parameter values for the various Tel1 models, we see that the 1D models would still be the best models for Tel1 even when constrained to have the same ChIP parameter values as Mec1.

## Supplemental Figures

**Figure S1:**
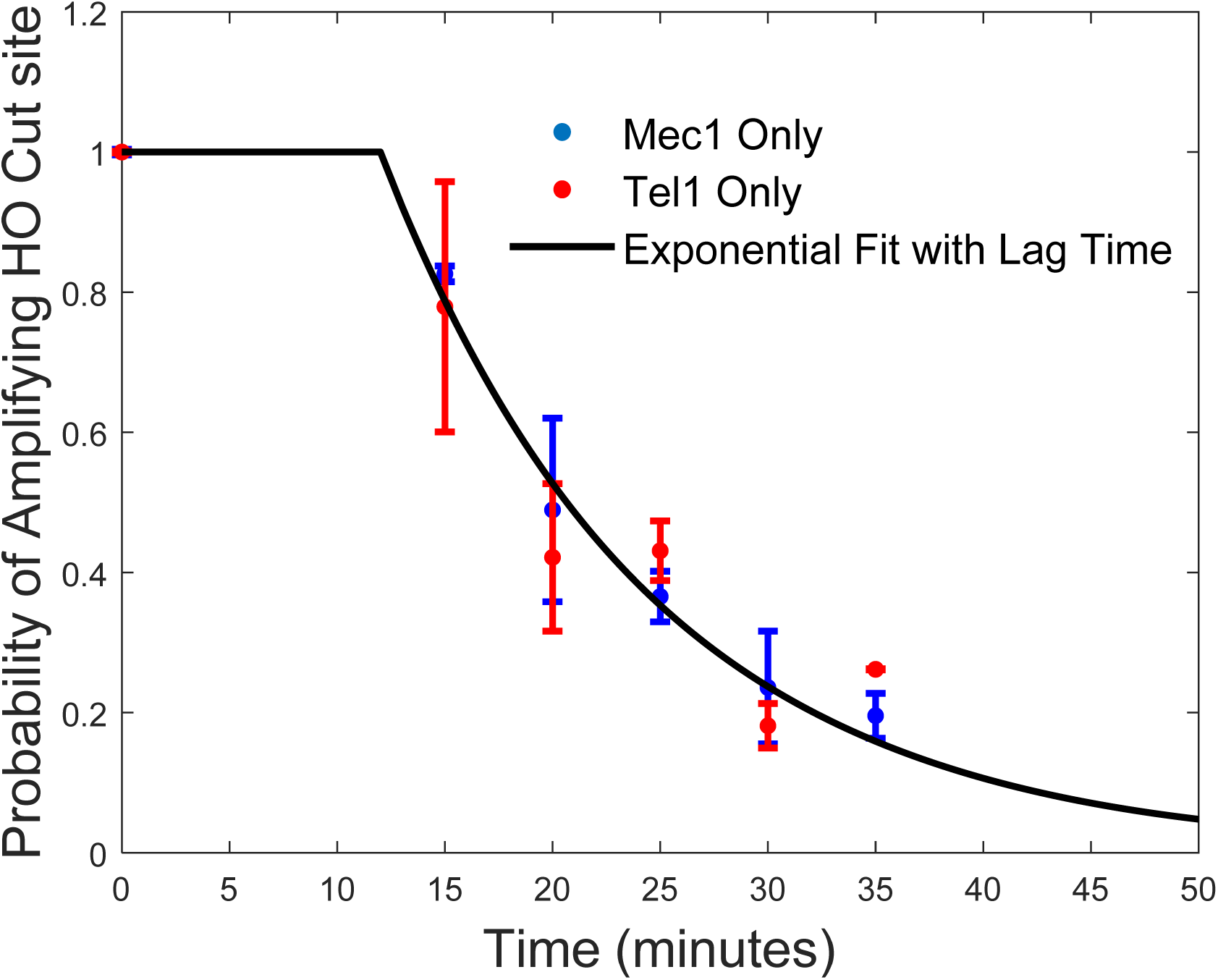
Kinetics of DSB induction. The amount of cleavage at the MAT locus by HO endonuclease after galactose induction was monitored in strains deleted for *YKU80* and carrying either Mec1 (blue) or Tel1 (red). qPCR was performed with primers flanking the cut site to quantify the levels of cleavage over 35 mins. In the presence of a DSB, qPCR fails to amplify the DNA. Approximately, 80% cutting was achieved by 30 mins. The black line is an exponential fit to both strains, using a lag time of 12 minutes (reflecting transcription and translation of the HO endonuclease) and a DNA cleavage rate of *k*_*DSB*_ = 0.08/minute. The lag time and *k*_*DSB*_ are used in calculating our models of *γ*-H2AX spreading. Error bars represent the standard error of the mean from n≥3 measurements.

**Figure S2:**
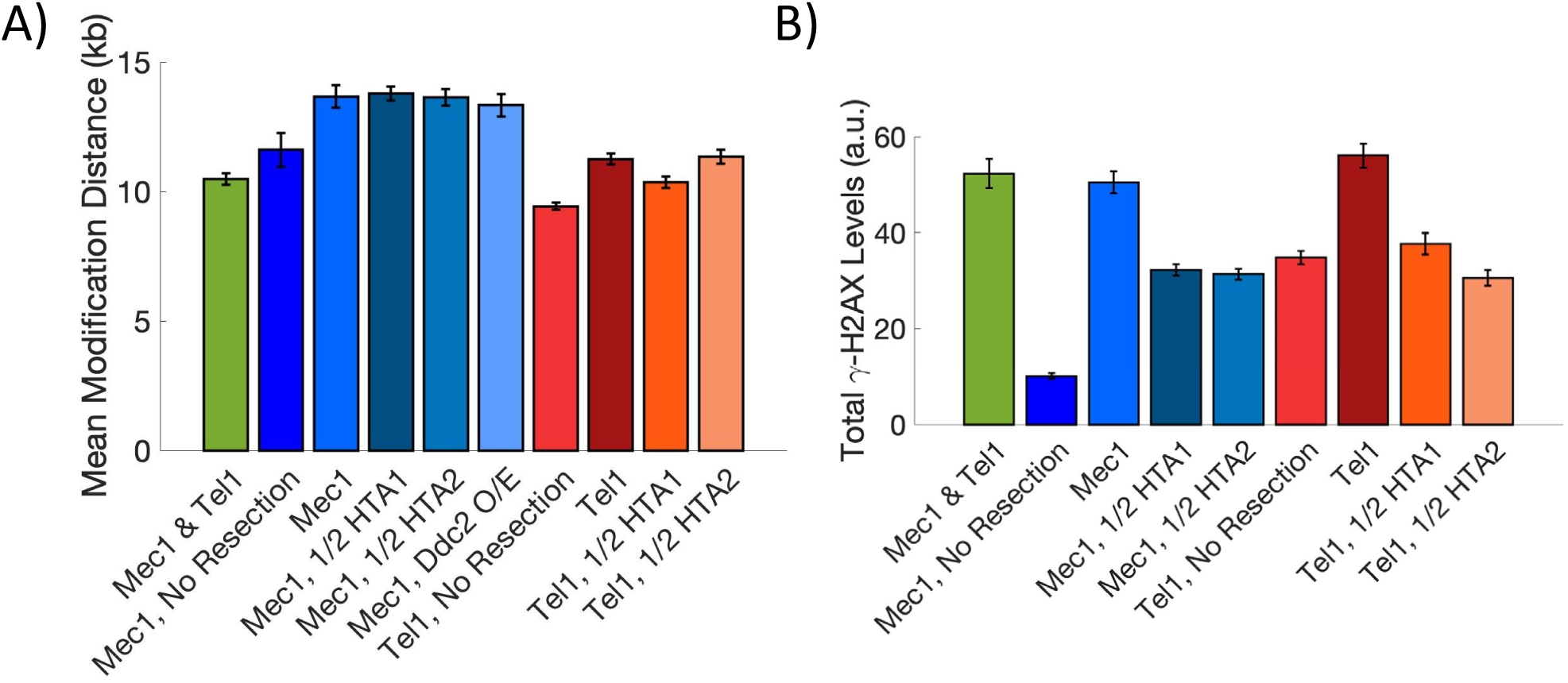
Mean modification distance and total γ-H2AX accumulation. **A)** The mean modification distance (MMD), the distance from the break that encompasses half the γ-H2AX profile, is displayed for all strains at 75 min except for Mec1, Ddc2 O/E which was measured at 60 min. Error bars represent the standard error of the mean. **B)** Total γ-H2AX levels were calculated by summing up γ-H2AX levels across all measured distances at 75 min. Error bars represent the standard error of the mean.

**Figure S3:**
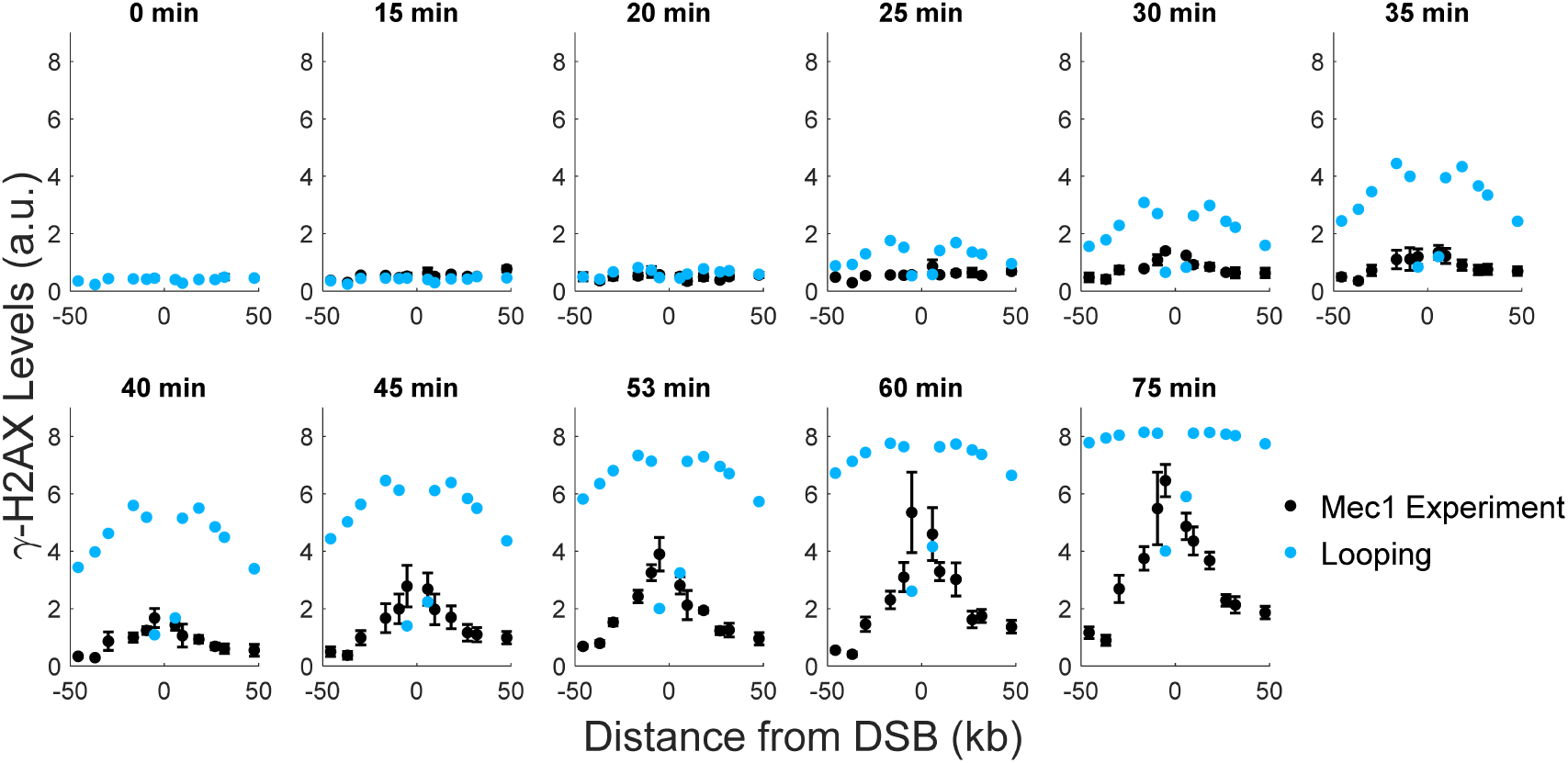
Comparison of experimental data by Mec1 to a looping model with a linearly increasing phosphorylation rate. Comparisons of the predicted phosphorylation levels to experimentally measured γ-H2AX levels. This model fits the data much worse than the best model, which is shown in Figure 5. Experimental error bars are the standard error of the mean from n≥3 measurements. Bayesian parameter estimation was used to simultaneously fit the model to the data shown here and to the RE-adjacent measurements from Figure 3. The optimal parameter values from Bayesian parameter estimation were used to plot the model, and are as follows: *C* = 44, *f* = 0.035, *N*_*H2A*_ = 6, *l* = 8.4 kb, and *ψ* = 0.039/minute^2^. In Figure 5 and Table 1B, Mec1 and Tel1 are constrained to the same values *N*_*H2A*_, *f* and *C* for the ChIP parameters.

**Figure S4:**
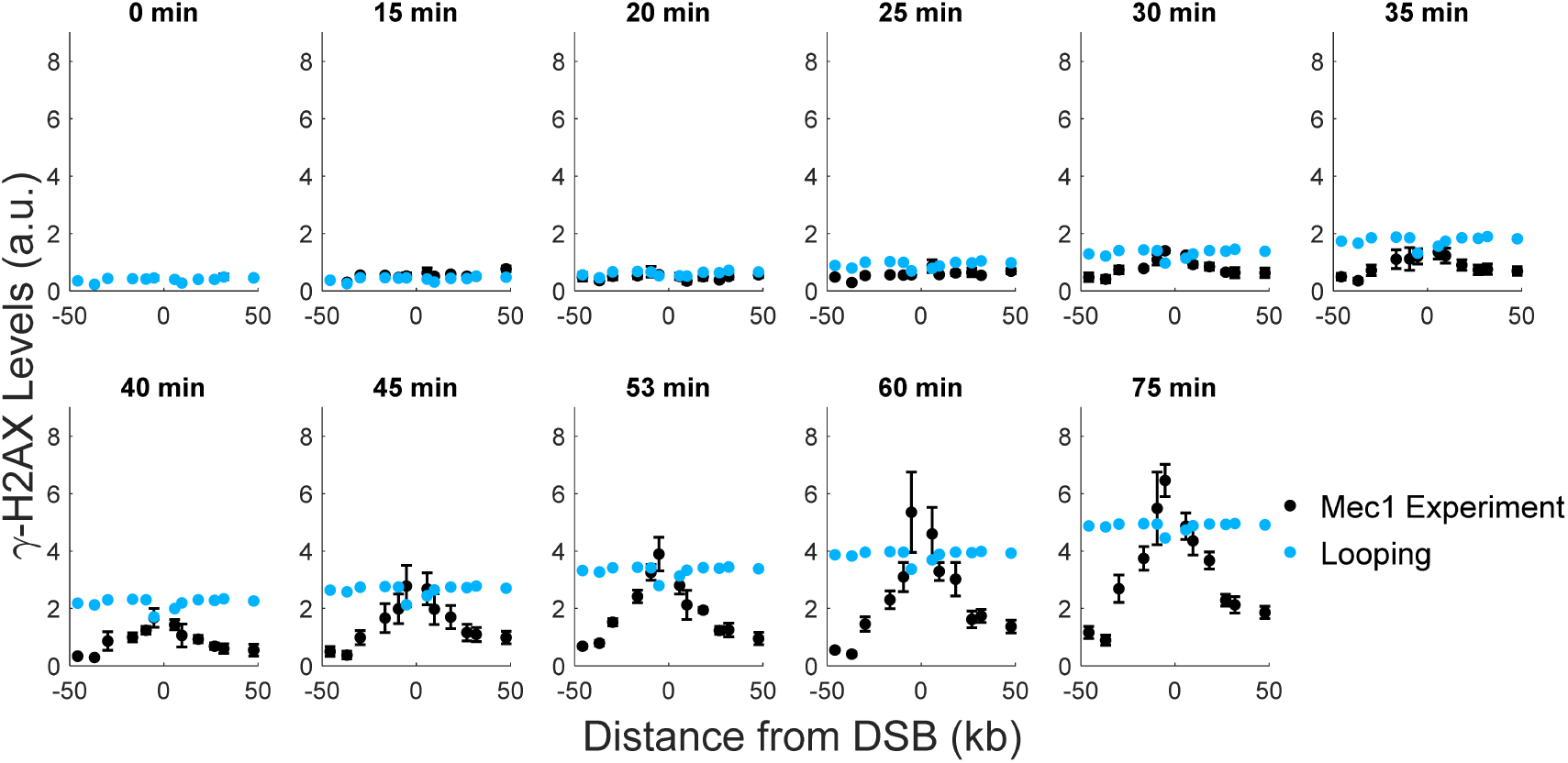
Comparison of experimental data by Mec1 to a looping model with a constant phosphorylation rate. The best model parameters are *C* = 120, *f* = 0.035, *N*_*H2A*_ = 2, *l* = 8.4 kb, *φ* = 11/minute, and *k*_*init*_ = 0.017/minute. For further details, refer to the caption of Figure S3.

**Figure S5:**
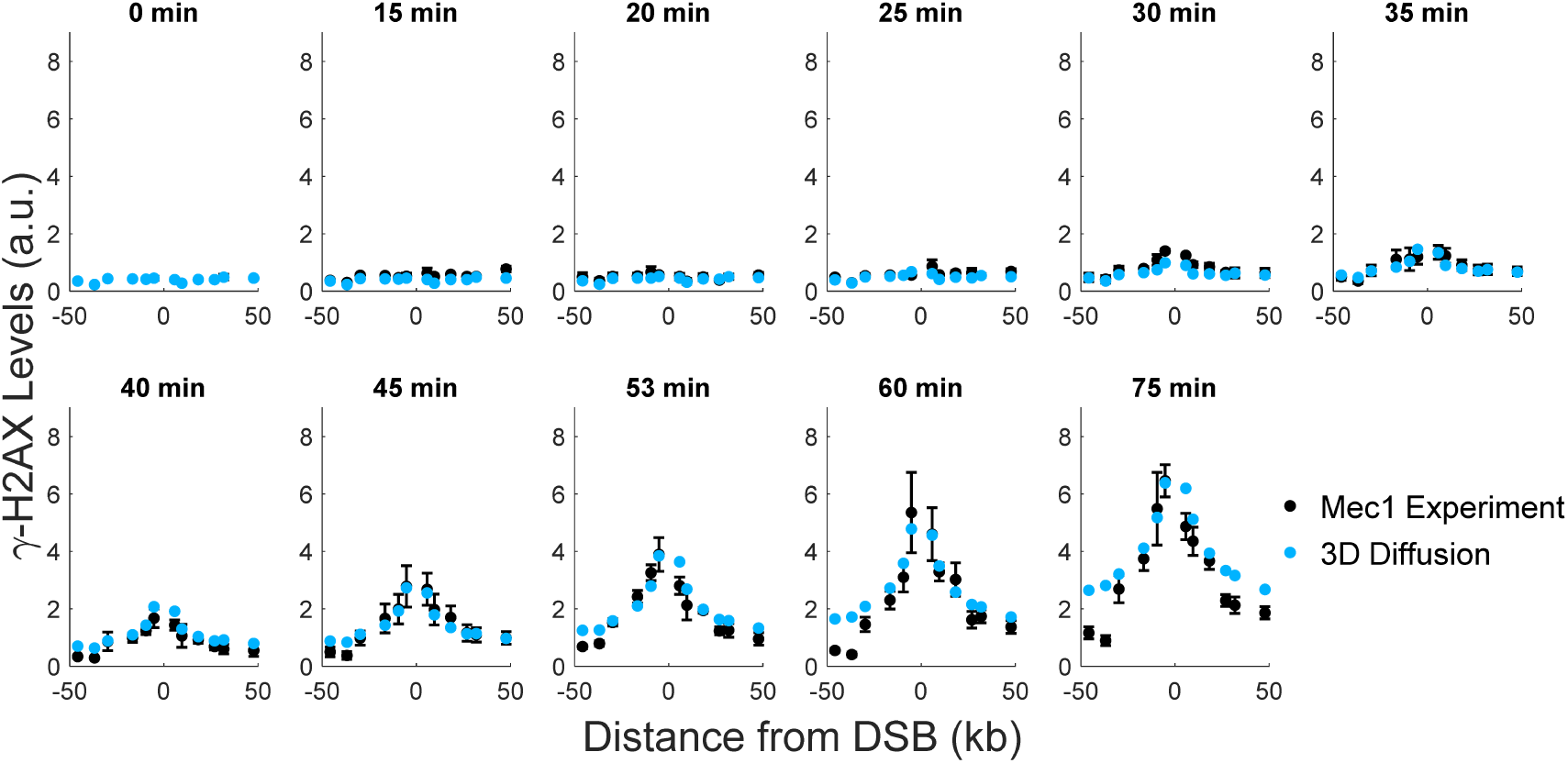
Comparison of experimental data by Mec1 to a 3D diffusion model with a linearly increasing rate of diffusion initiation. The best model parameters are *C* = 13, *f* = 0.16, *N*_*H2A*_ = 6, *l* = 15 kb, and *ζ* = 3.7×10^−4^/minute^2^. For further details, refer to the caption of Figure S3.

**Figure S6:**
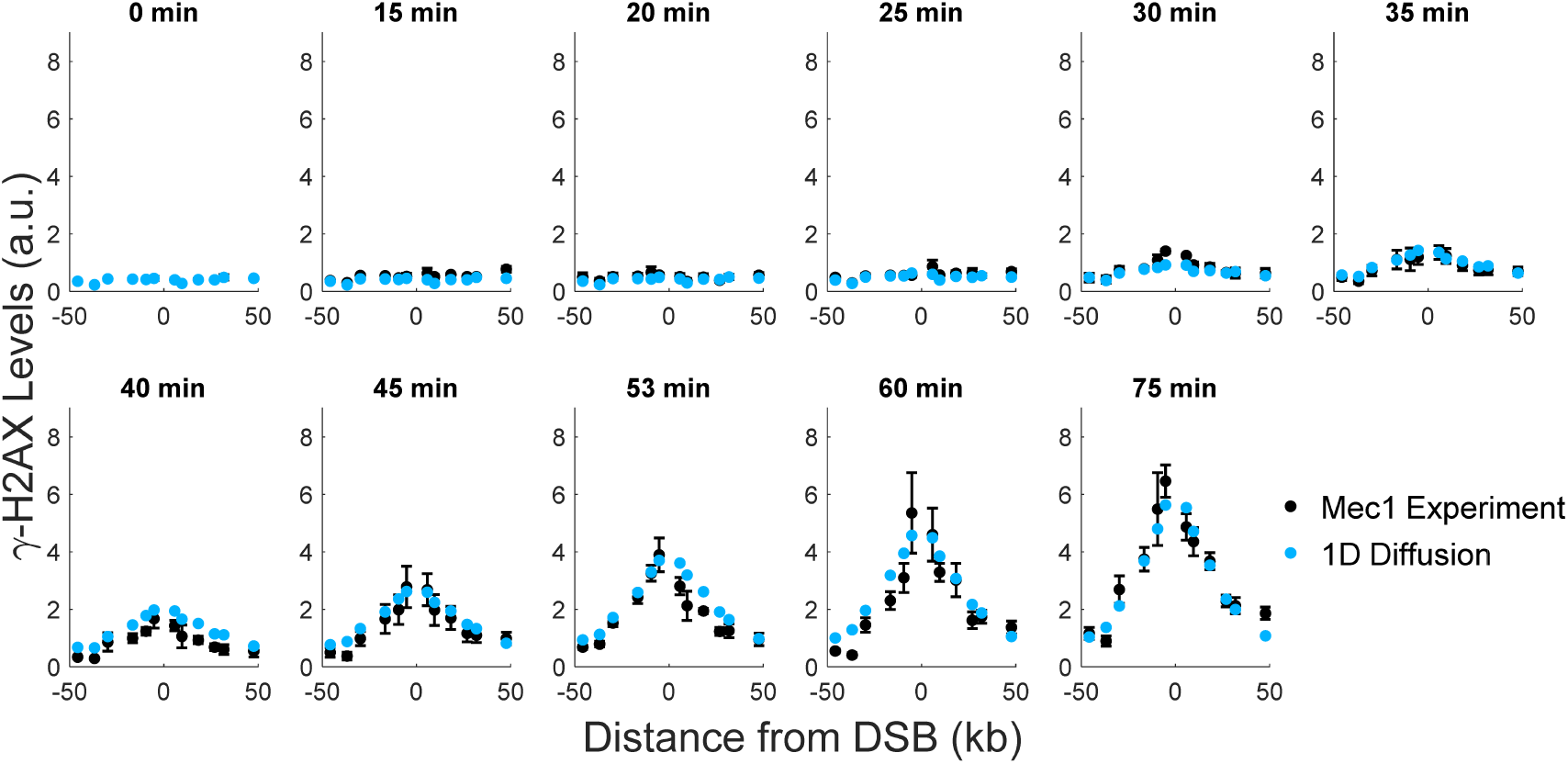
Comparison of experimental data by Mec1 to a 1D diffusion model with a linearly increasing rate of diffusion initiation. The best model parameters are *C* = 11, *f* = 0.80, *N*_*H2A*_ = 6, *D* = 18 kb^2^/minute, *k*_*cat*_ = 0.039/minute, and *z* = 84/minute^2^. For further details, refer to the caption of Figure S3.

**Figure S7:**
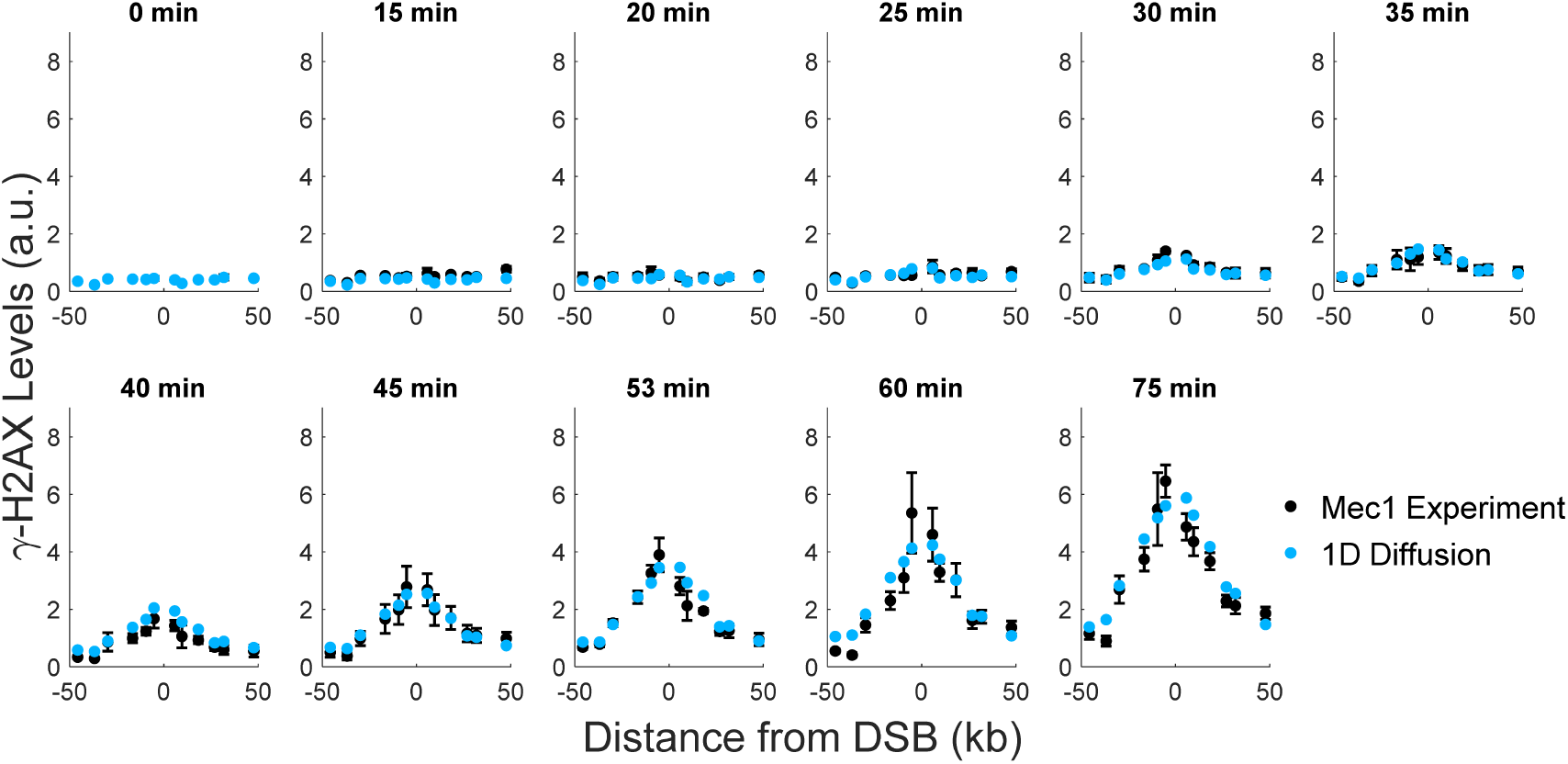
Comparison of experimental data by Mec1 to a 1D diffusion model with a constant rate of diffusion initiation. The best model parameters are *C* = 32, *f* = 0.28, *N*_*H2A*_ = 6, *D* = 22 kb^2^/minute, *k*_*cat*_ = 1.9/minute, and *k*_*init*_ = 0.019/minute. For further details, refer to the caption of Figure S3.

**Figure S8:**
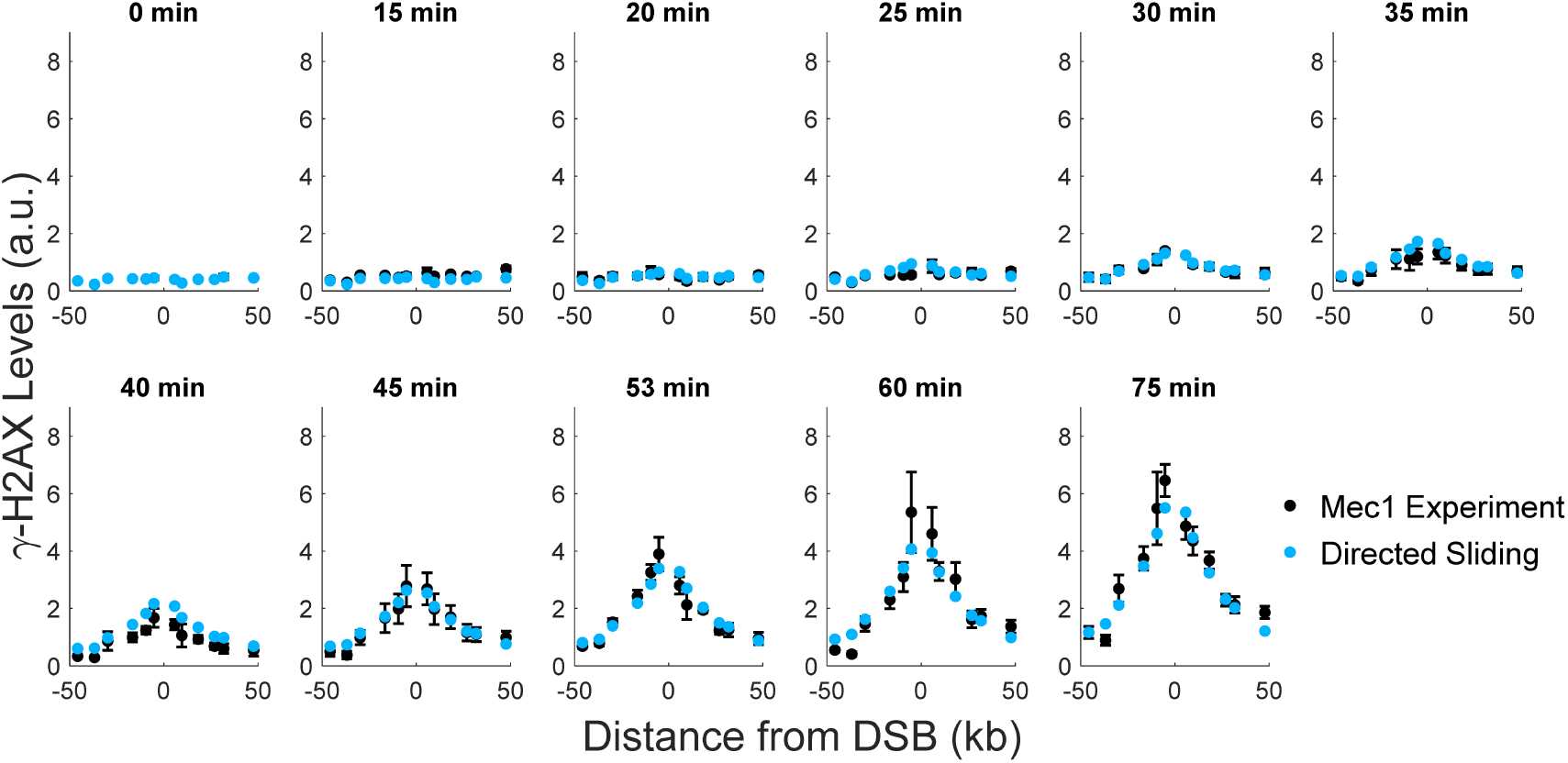
Comparison of experimental by Mec1 to a directed sliding model. The best model parameters are *C* = 73, *f* = 0.72, *N*_*H2A*_ = 2, *k*_*slide*_ = 17 kb/minute, *k*_*off*_ = 0.74/minute, and *k*_*init*_ = 0.0020/minute. For further details, refer to the caption of Figure S3.

**Figure S9:**
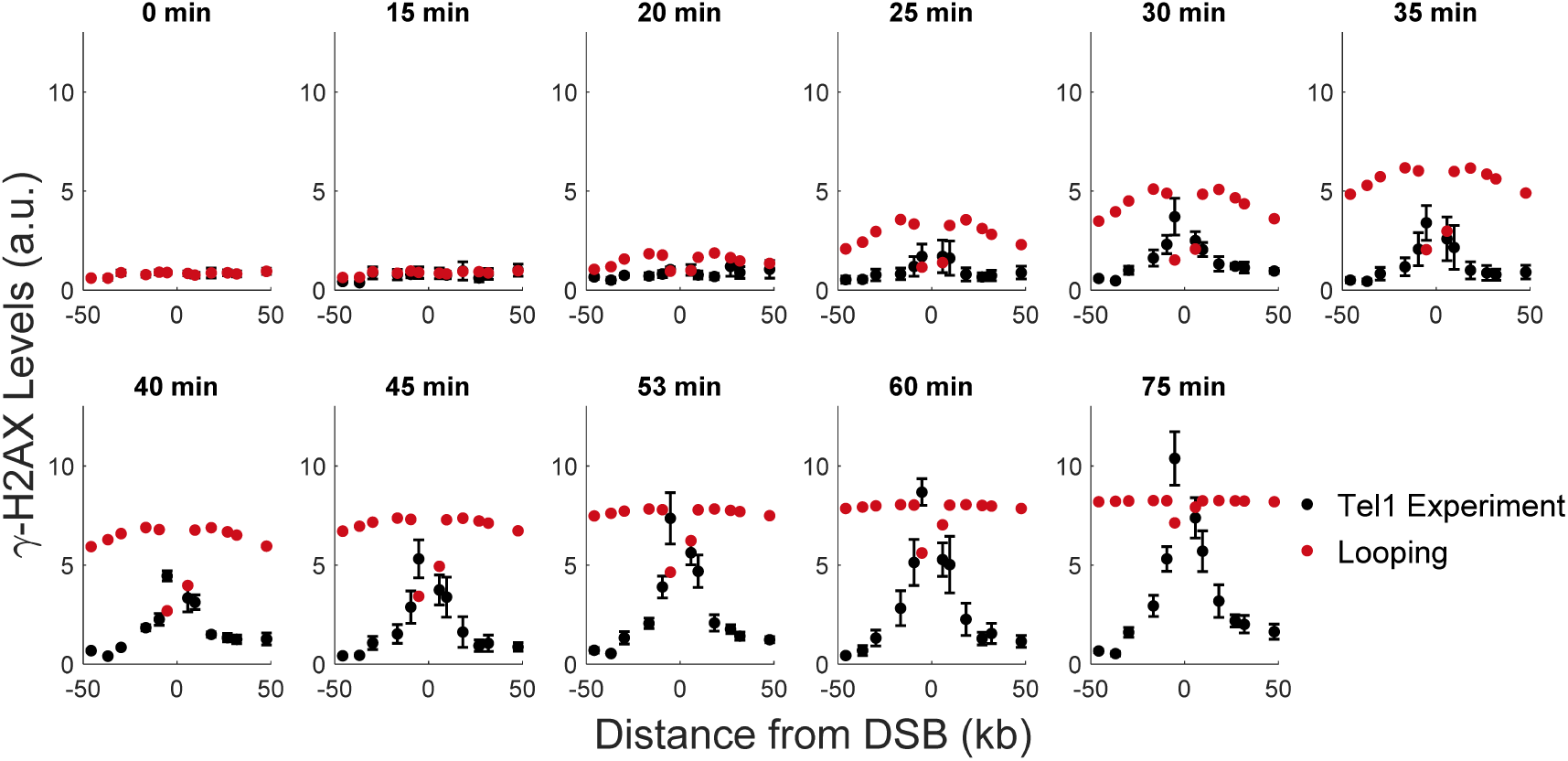
Comparison of experimental data by Tel1 to a looping model with a linearly increasing phosphorylation rate. The best model parameters are *C* = 120, *f* = 0.035, *N*_*H2A*_ = 2, *l* = 8.4 kb, and *ψ* = 0.14/minute^2^. For further details, refer to the caption of Figure S3.

**Figure S10:**
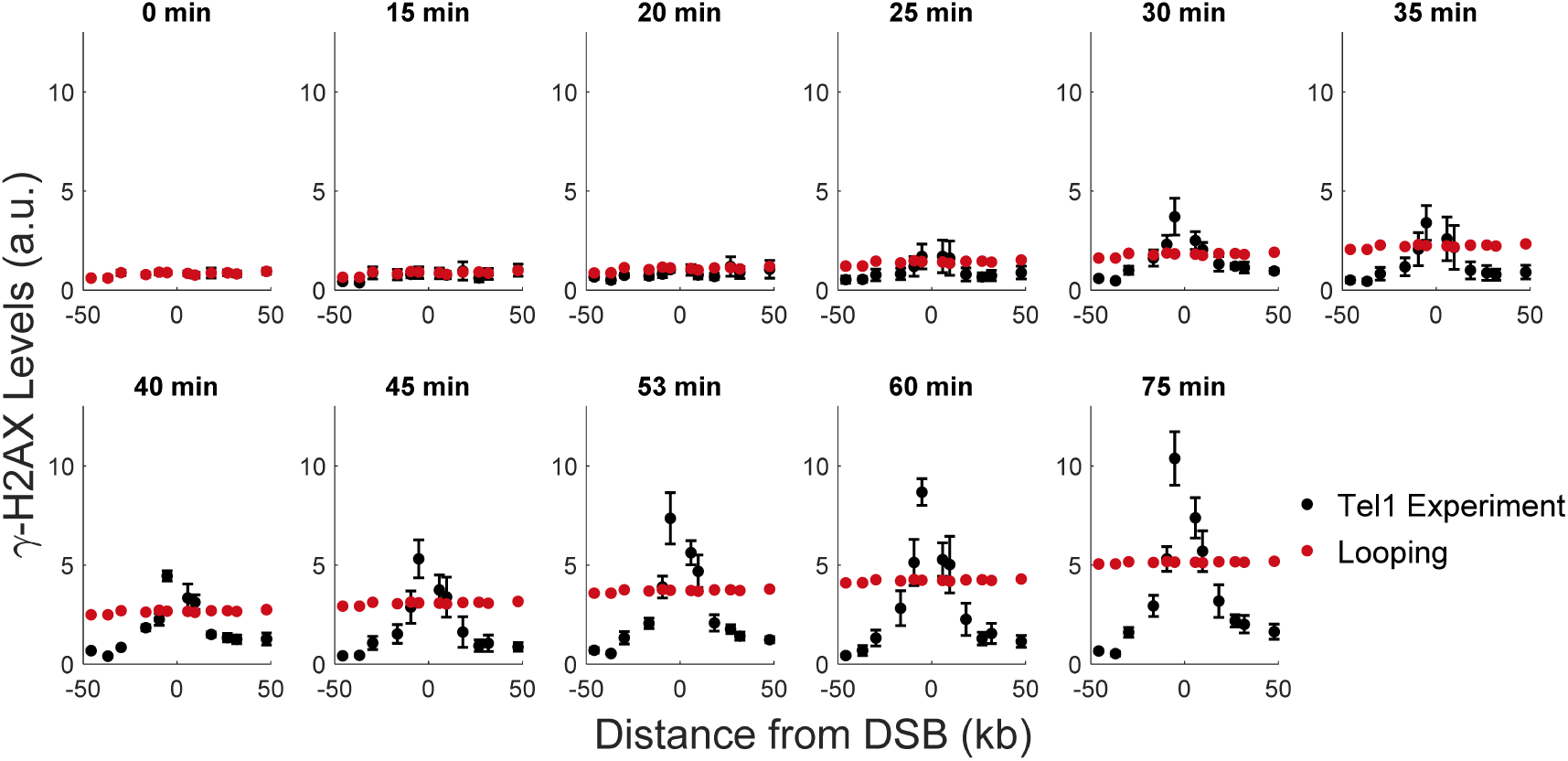
Comparison of experimental data by Tel1 to a looping model with a constant phosphorylation rate. The best model parameters are *C* = 120, *f* = 0.035, *N*_*H2A*_ = 2, *l* = 8.4 kb, *φ* = 200/minute, and *k*_*init*_ = 0.017/minute. For further details, refer to the caption of Figure S3.

**Figure S11:**
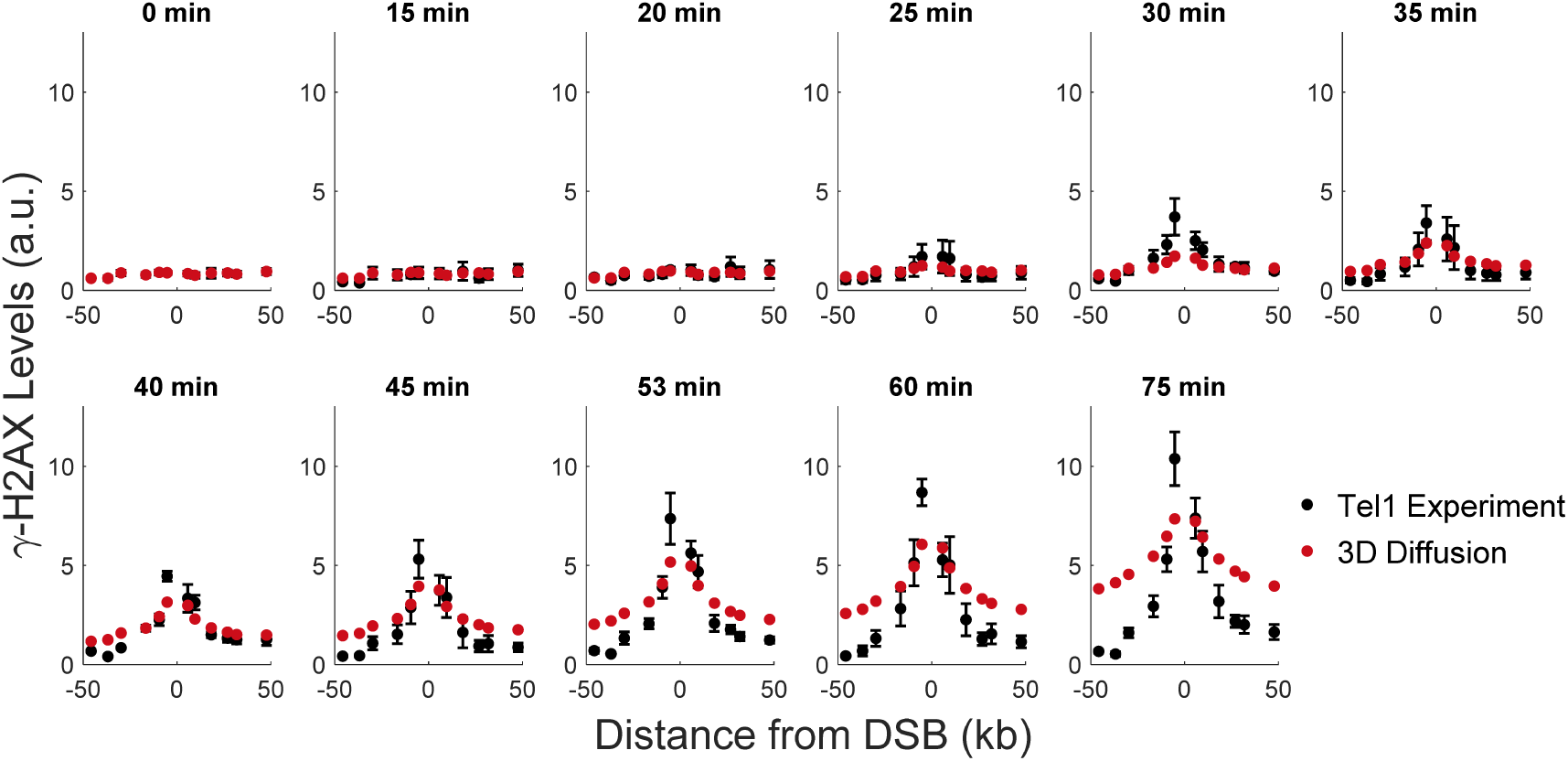
Comparison of experimental by Tel1 to a 3D diffusion model with a linearly increasing rate of diffusion initiation. The best model parameters are *C* = 13, *f* = 0.16, *N*_*H2A*_ = 6, *l* = 15 kb, and *ζ* = 6.2×10^−4^/minute^2^. For further details, refer to the caption of Figure S3.

**Figure S12:**
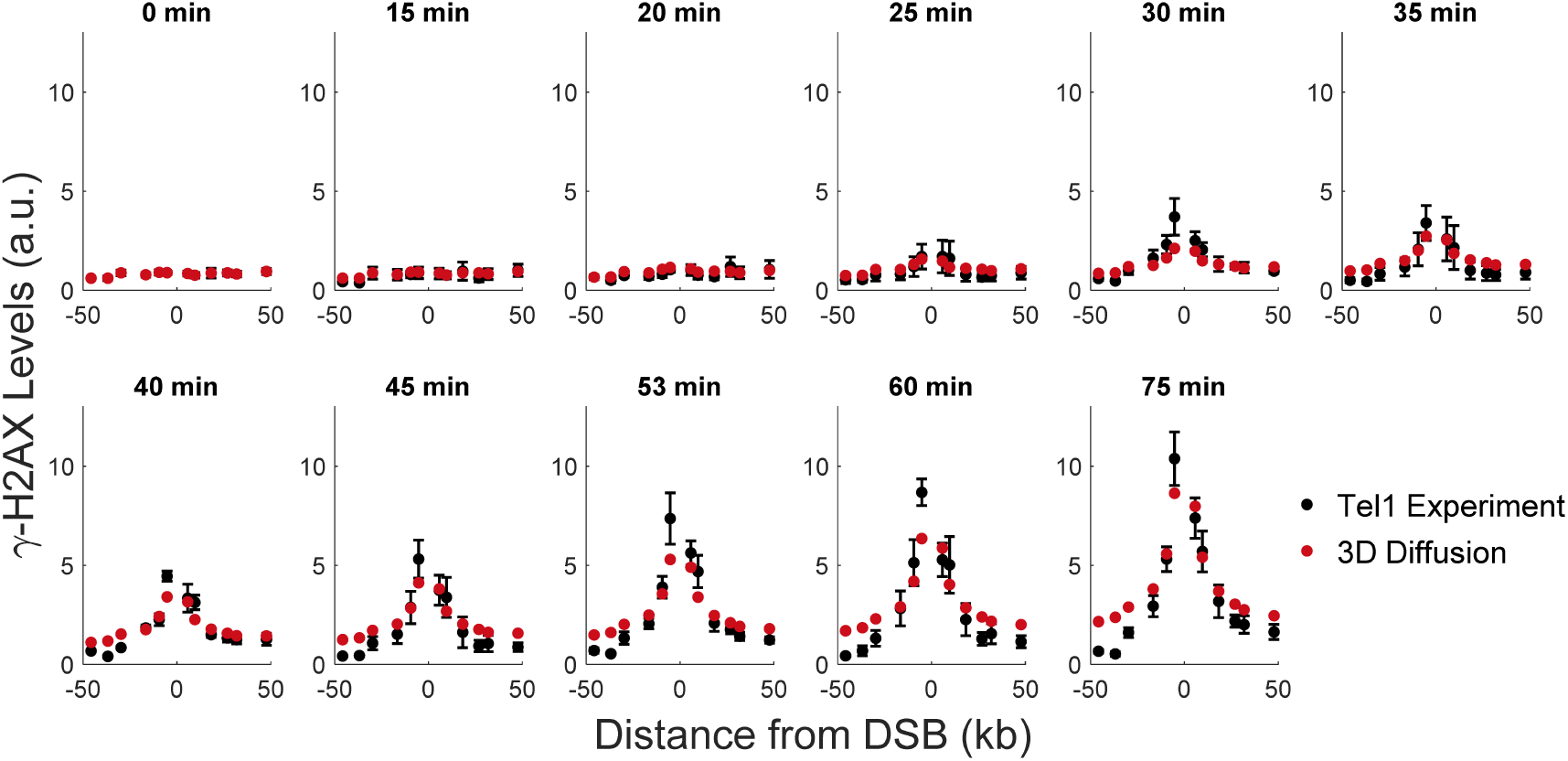
Comparison of experimental data by Tel1 to a 3D diffusion model with a constant rate of diffusion initiation. The best model parameters are *C* = 73, *f* = 0.55, *N*_*H2A*_ = 6, *l* = 15 kb, *ω* = 0.00030/minute, and *k*_*init*_ = 1.0/minute. For further details, refer to the caption of Figure S3.

**Figure S13:**
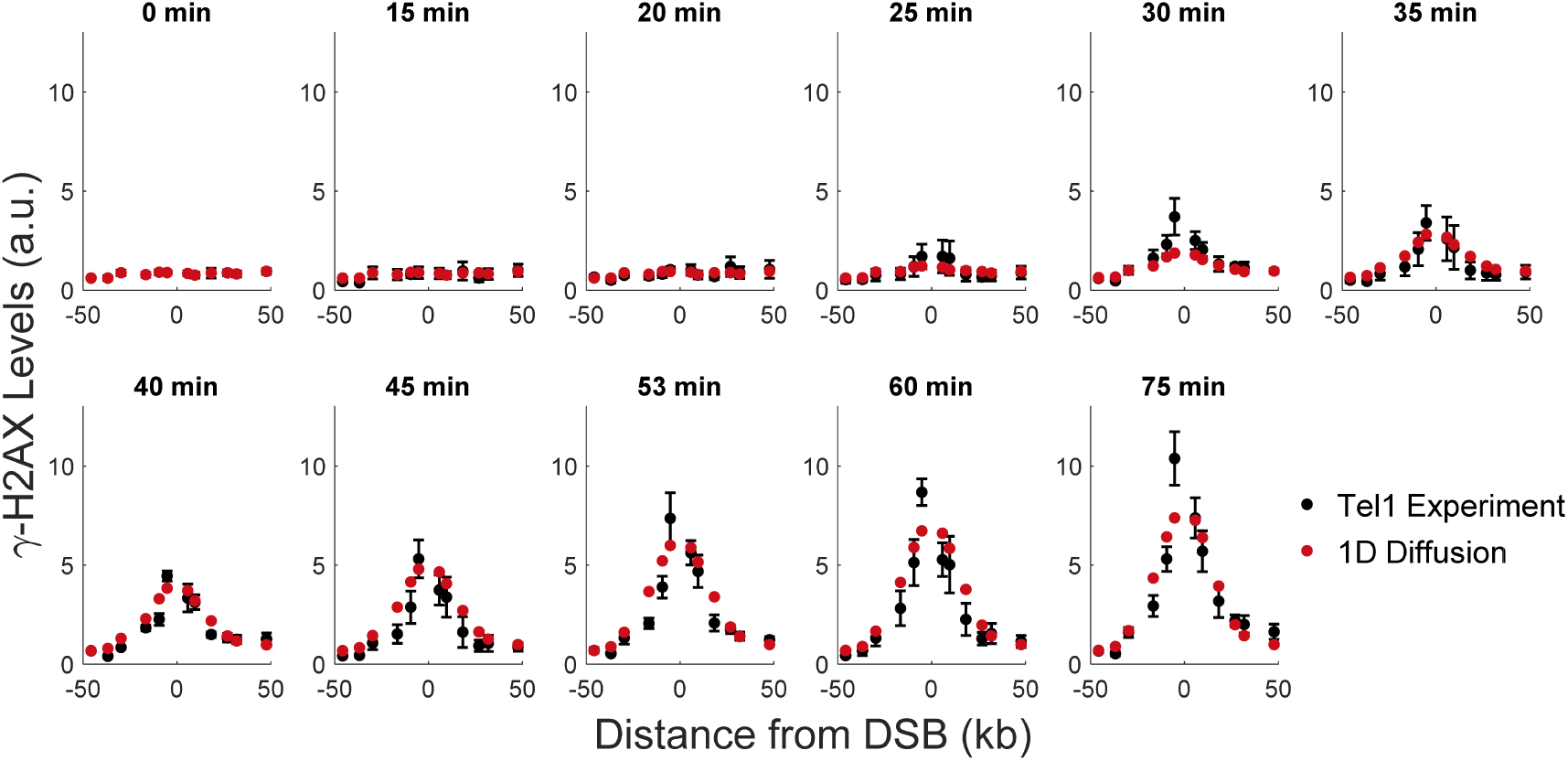
Comparison of experimental data by Tel1 to a 1D diffusion model with a linearly increasing rate of diffusion initiation. The best model parameters are *C* = 9.3, *f* = 0.50, *N*_*H2A*_ = 4, *D* = 3.9 kb^2^/minute, *k*_*cat*_ = 0.59/minute, and *z* = 37/minute^2^. For further details, refer to the caption of Figure S3.

**Figure S14:**
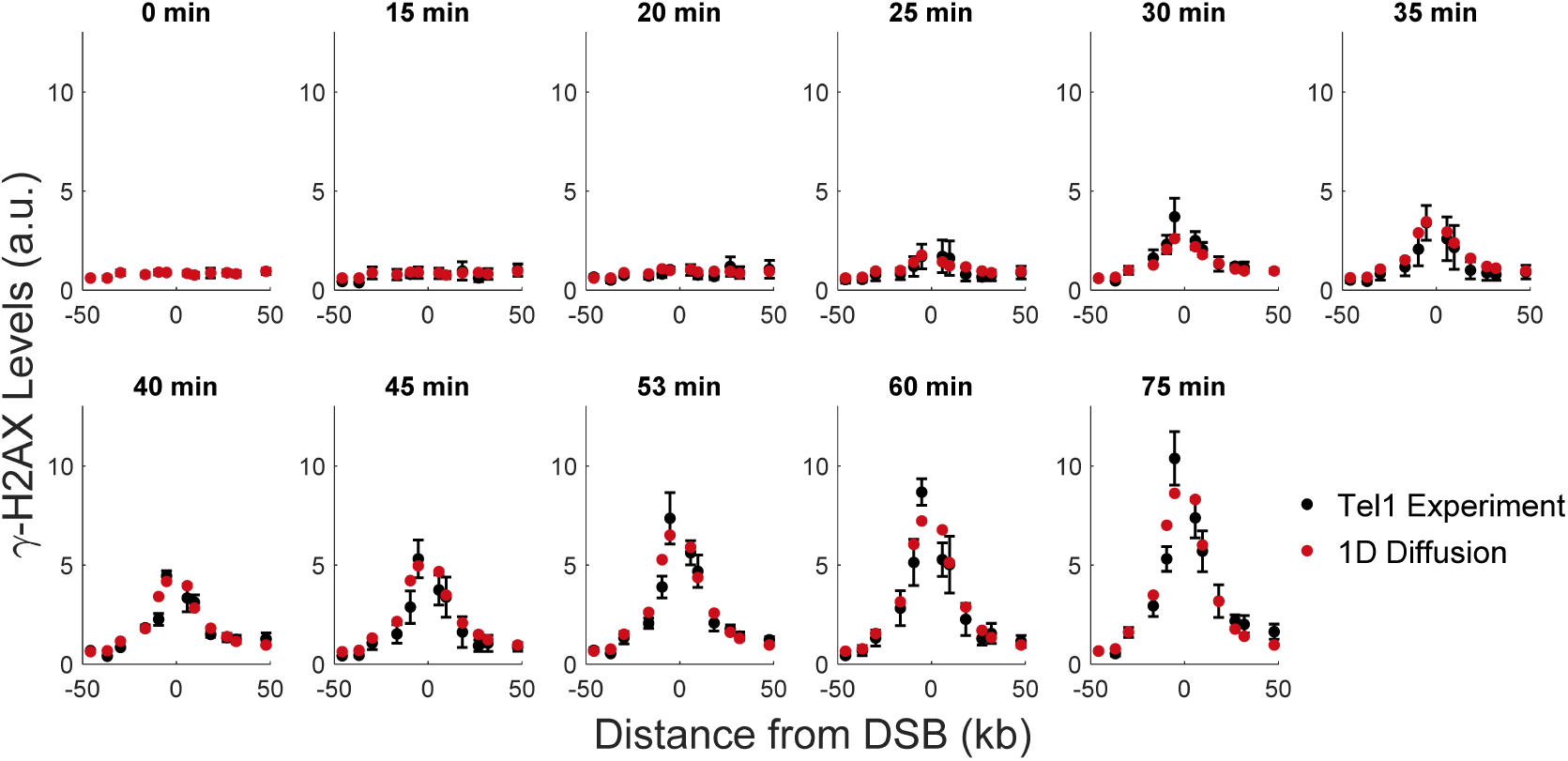
Comparison of experimental data by Tel1 to a 1D diffusion model with a constant rate of diffusion initiation. The best model parameters are *C* = 32, *f* = 0.55, *N*_*H2A*_ = 4, *D* = 4.1 kb^2^/minute, *k*_*cat*_ = 0.20/minute, and *k*_*init*_ = 0.098/minute. For further details, refer to the caption of Figure S3.

## Supplemental Tables

**Table S1:** log_10_(Bayes Factor) for all mechanisms. The log_10_(Bayes Factor) is shown for every model variant. (Table 1 showed the Bayes factors for models with a constant rate of diffusion initiation or constant phosphorylation rate). The Bayes factor is calculated by dividing the probability of the indicated model by the probability of the best model. Bayes factors were computed for Mec1 and Tel1 separately. For Mec1, the best model is 3D diffusion with a constant rate of diffusion initiation. For Tel1, the best model is directed sliding.

| Model | Mec1, $\log_{10}$(Bayes Factor) | Tel1, $\log_{10}$(Bayes Factor) |
| --- | --- | --- |
| Looping with Linearly Increasing Phosphorylation Rate | -545 | -449 |
| Looping with Constant Phosphorylation Rate | -252 | -251 |
| 3D Diffusion with Linearly Increasing Rate of Diffusion Initiation | -28 | -136 |
| 3D Diffusion with Constant Rate of Diffusion Initiation | Best Model | -61 |
| Directed Sliding | -14 | Best Model |
| 1D Diffusion with Linearly Increasing Rate of Diffusion Initiation | -28 | -35 |
| 1D Diffusion with Constant Rate of Diffusion Initiation | -12 | -5 |

**Table S2:**
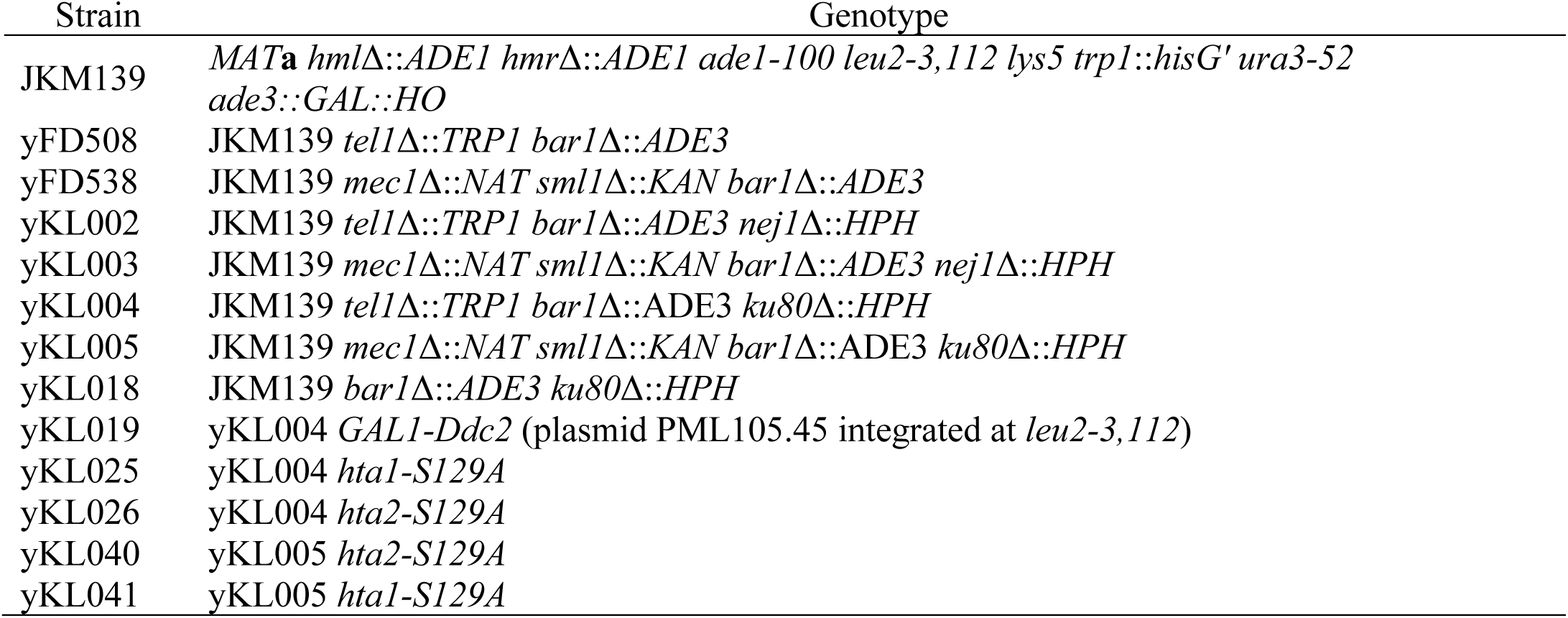
List of yeast strains used in this study.

**Table S3:**
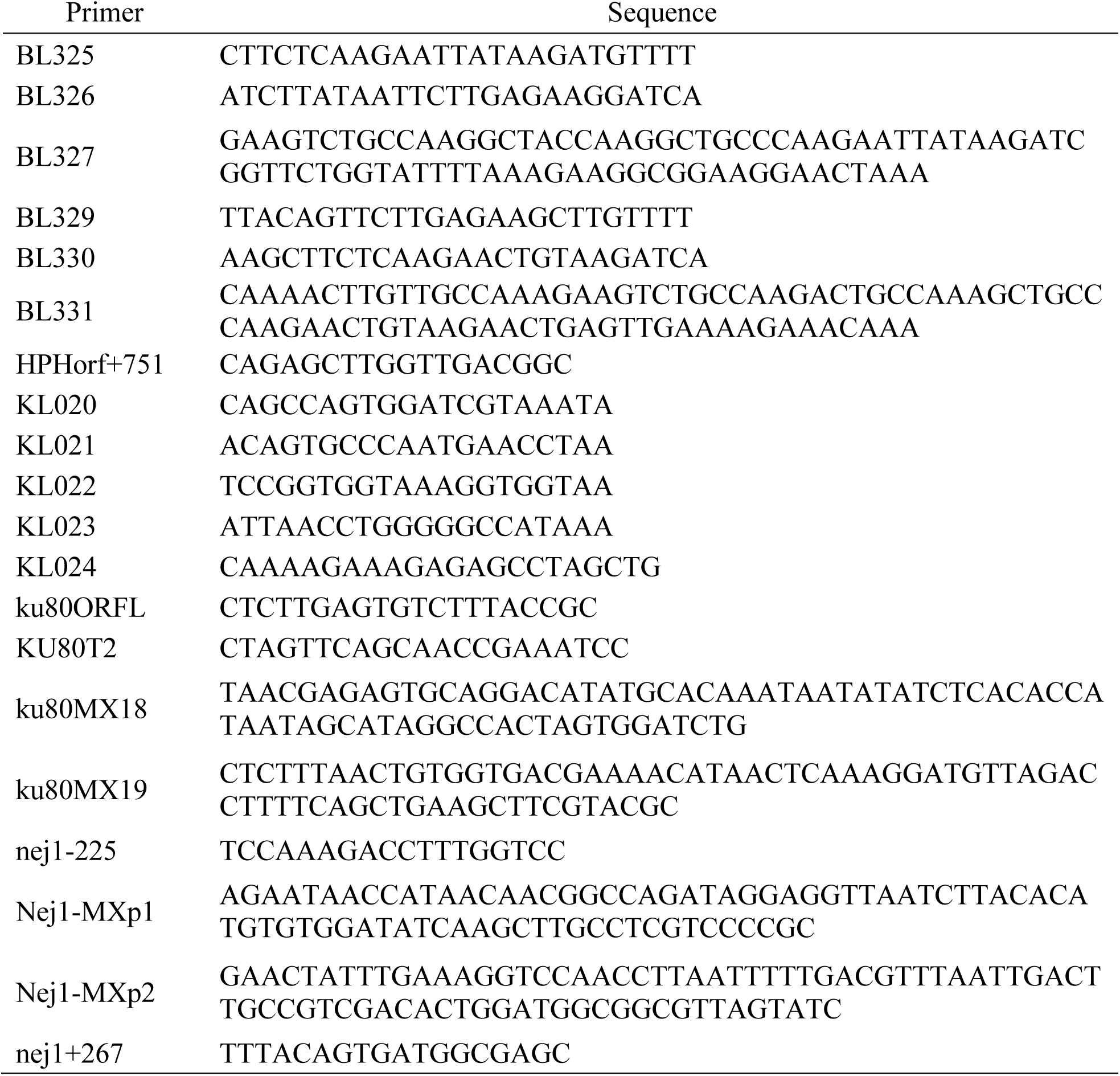
Primers used during strain construction.

**Table S4:** Plasmids used during strain construction.

| Plasmid | Genotype |
| --- | --- |
| pAG32 | ampr, *HPH* |
| pBL13 | ampr, *Cas9*, *URA3*, gRNA targeting *HTA1* |
| pBL14 | ampr, *Cas9*, *URA3*, gRNA targeting *HTA2* |
| pJH2972 | ampr, *Cas9*, *URA3* |
| PML105.45 | *GAL1-Ddc2*, *LEU2* |

**Table S5:**
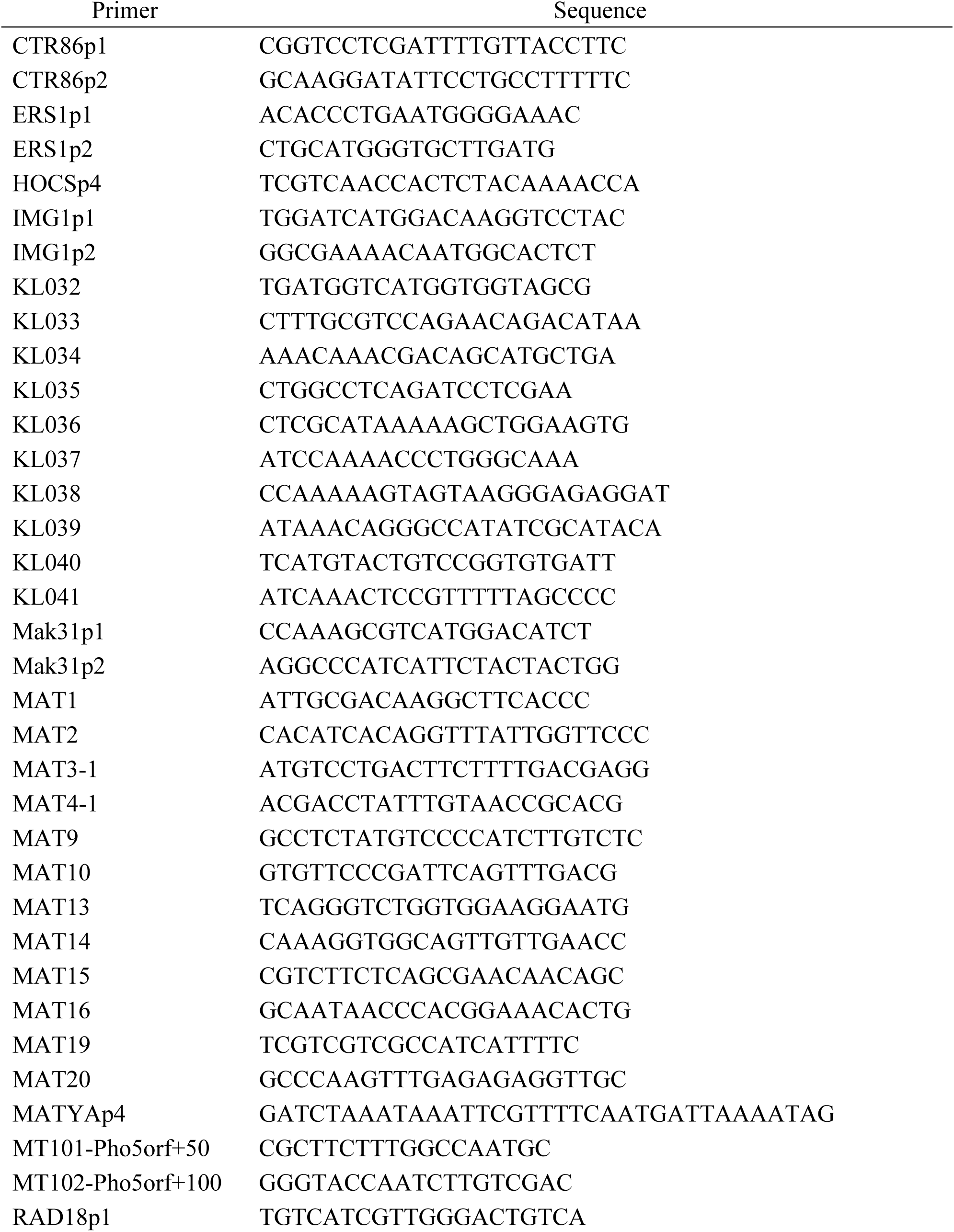
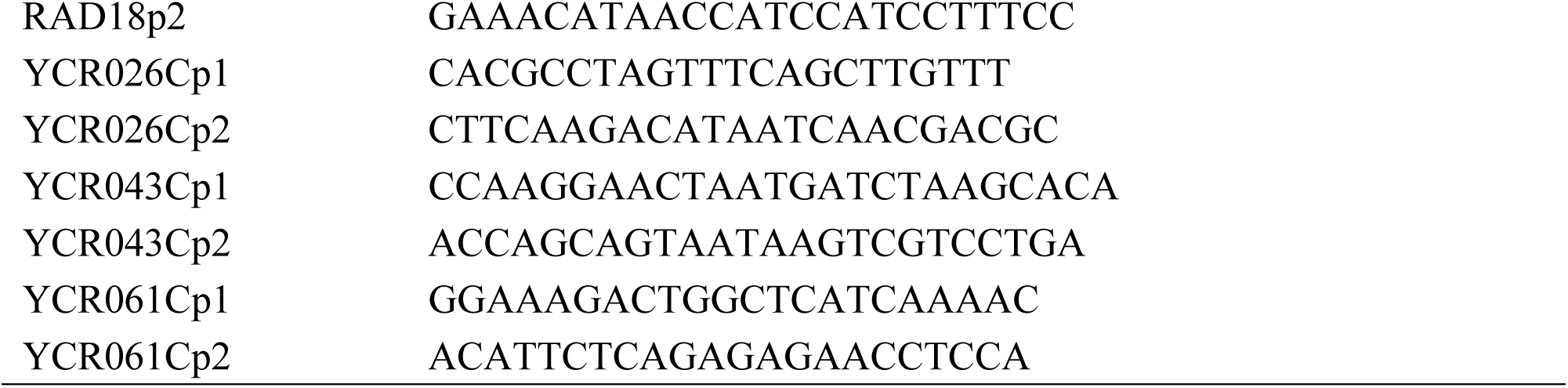
Primers used in quantitative PCR (qPCR)

## Supplemental Dataset

### Dataset S1: γ-H2AX levels around MAT and RE

γ-H2AX levels are shown in the Excel file “Dataset S1.” γ-H2AX levels were measured around the MAT locus for strains listed in Table S2. γ-H2AX measurements around RE were measured for strains yKL004 and yKL005.

